# An octopamine-specific GRAB sensor reveals a monoamine relay circuitry that boosts aversive learning

**DOI:** 10.1101/2024.03.09.584200

**Authors:** Mingyue Lv, Ruyi Cai, Renzimo Zhang, Xiju Xia, Xuelin Li, Yipan Wang, Huan Wang, Jianzhi Zeng, Yifei Xue, Lanqun Mao, Yulong Li

## Abstract

Octopamine (OA), analogous to norepinephrine in vertebrates, is an essential monoamine neurotransmitter in invertebrates that plays a significant role in various biological functions, including olfactory associative learning. However, the spatial and temporal dynamics of OA *in vivo* remain poorly understood due to limitations associated with the currently available methods used to detect it. To overcome these limitations, we developed a genetically encoded <u>G</u>PC<u>R</u> <u>a</u>ctivation-<u>b</u>ased (GRAB) OA sensor called GRAB_OA1.0_. This sensor is highly selective for OA and exhibits a robust and rapid increase in fluorescence in response to extracellular OA. Using GRAB_OA1.0_, we monitored OA release in the *Drosophila* mushroom body (MB), the fly’s learning center, and found that OA is released in response to both odor and shock stimuli in an aversive learning model. This OA release requires acetylcholine (ACh) released from Kenyon cells, signaling via nicotinic ACh receptors. Finally, we discovered that OA amplifies aversive learning behavior by augmenting dopamine-mediated punishment signals via Octβ1R in dopaminergic neurons, leading to alterations in synaptic plasticity within the MB. Thus, our new GRAB_OA1.0_ sensor can be used to monitor OA release in real-time under physiological conditions, providing valuable insights into the cellular and circuit mechanisms that underlie OA signaling.

## INTRODUCTION

Octopamine (OA) is an essential monoamine neurotransmitter in invertebrates, analogous to norepinephrine (NE) in vertebrates[1, 2]. In vertebrates, OA is classified as a trace amine and is thought to be associated with emotional responses[3–5]. In invertebrates, OA plays a role in various physiological processes, including the sleep-wake cycle, flight, ovulation, aggression, and associative learning[6–27].

In *Drosophila melanogaster*, OA has been implicated in regulating both learning and memory, particularly in the formation of short-term associative memories of an odor-conditioned stimulus (CS) paired with either an appetitive sugar reward or an aversive electrical body shock as the unconditioned stimulus (US). Moreover, studies have shown that mutants lacking tyramine β hydroxylase (TβH), the rate-limiting enzyme for OA biosynthesis, have an impaired ability to acquire appetitive memory[19]. Furthermore, stimulation of octopaminergic neurons (OANs) can replace sugar presentation during conditioning and lead to the formation of short-term appetitive memory[20, 21]. However, studies regarding aversive conditioning have yielded conflicting results. For example, some studies found normal performance in TβH mutants[19, 28], while other studies found impaired performance when compared to wild-type (WT) flies[29].

In the *Drosophila* brain, the mushroom body (MB) is the main center for olfactory learning[30–33] and consists primarily of Kenyon cells (KCs), with their dendrites residing in the calyx and their axon bundles projecting through the peduncle to form the α/β lobe, α’/β’ lobe and γ lobe[34–36]. Studies have shown that OA signaling via the β-adrenergic-like OA receptor Octβ1R is required for aversive memory formation in the MB[25]. In addition to its role in short-term memory, OA released from the anterior paired lateral (APL) neurons has been shown to modulate intermediate-term aversive memory by acting on KCs via Octβ2R[23]. Together, these findings suggest that OA indeed plays a key role in aversive learning and memory in *Drosophila*. However, there are still many unresolved issues regarding the spatiotemporal dynamics of OA release and the specific role OA plays in aversive learning that warrant further investigations.

Our relatively limited understanding of how OA functions spatially and temporally during learning is primarily due to limitations in current detection methods. Traditional methods, such as microdialysis-coupled biochemical analysis[37–39], offer high specificity but low temporal resolution and complex sampling procedures, especially in invertebrates. On the other hand, electrochemical techniques like fast-scan cyclic voltammetry (FSCV) enable rapid monitoring of endogenous OA release[40, 41], but they cannot distinguish between OA and other structurally similar neurotransmitters, particularly its biological precursor tyramine (TA), which differs from OA by only one hydroxyl group and also serves as an important monoamine in invertebrates[2].

To overcome these limitations, we developed a novel G protein-coupled receptor (GPCR) activation-based (GRAB) OA sensor, utilizing the *Drosophila* Octβ2R as the sensing module and circularly-permutated enhanced green fluorescent protein (cpEGFP) as the reporter; we call this sensor GRAB_OA1.0_ (hereafter referred to as OA1.0). We found that this sensor is highly specific to OA, has sub-second kinetics, and exhibits a peak increase in fluorescence of approximately 660% in response to OA. Using OA1.0, we then measured spatiotemporal changes of OA in the *Drosophila* MB in response to odor and shock stimuli. Our findings reveal that the release of OA in the MB promotes the release of dopamine (DA), which increases the fly’s perception of the US, thereby facilitating aversive learning.

## RESULTS

### Development and characterization of GRABOA1.0

To monitor octopamine (OA) release *in vivo* with high specificity, sensitivity and spatiotemporal resolution, we employed a well-established strategy[42–53] to develop a genetically encoded GPCR activation-based (GRAB) sensor for OA using EGFP to report an increase in extracellular OA through an increase in fluorescence intensity. First, we inserted the conformationally sensitive cpEGFP into the third intracellular loop (ICL3) of the β-adrenergic-like OA receptor Octβ2R. Next, we systematically screened the position of the cpEGFP and optimized the linker residues between the GPCR and cpEGFP using site-directed mutagenesis. Finally, we introduced mutations in the ligand-binding pocket of the GPCR to create the GRAB_OA1.0_ (OA1.0) sensor (Fig. 1A, B and Fig. S1).

**Figure 1.**
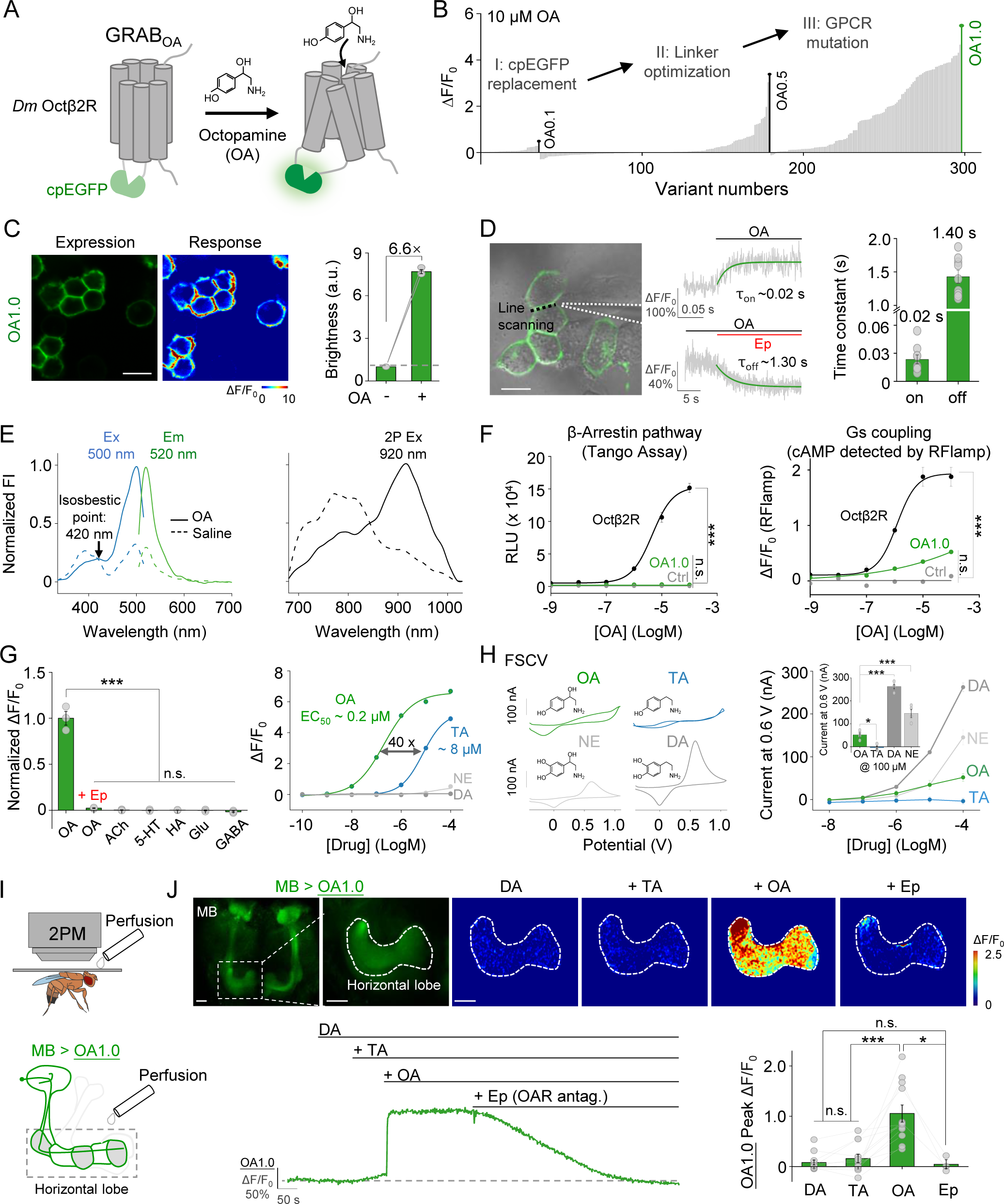
Development and characterization of the GRAB_OA1.0_ (OA1.0) sensor in HEK293T cells and living flies. (A) Schematic illustration depicting the strategy for developing the GRAB_OA_ sensor. Ligand binding activates the sensor, inducing a change in EGFP fluorescence. (B) Screening and optimization steps of GRAB_OA_ sensors, and the resulting change in fluorescence (ΔF/F_0_) in response to 10 μM OA. (C) Expression, fluorescence change in response to 100 μM OA, and summary data measured in HEK293T cells expressing OA1.0; n = 3 wells containing >500 cells each. (D) τ_on_ and τ_off_ were measured in OA1.0-expressing cells in response to OA and epinastine (Ep), respectively, in line-scan mode; an example image (left), representative traces (middle), and summary data (right) are shown; n ≥ 9 cells from 3 cultures; the dotted black line in the image indicates the line-scanning region. (E) One-photon (1P) excitation (ex) and emission (em) spectra (left) and two-photon (2P) excitation spectra (right) of OA1.0 were measured in the absence and presence of OA; FI, fluorescence intensity. (F) Left: The Tango assay was used to measure β-arrestin‒mediated signaling in cells expressing OA1.0 or wild-type (WT) Octβ2R and treated with increasing concentrations of OA; n = 3 wells containing >1000 cells each. Right: The RFlamp assay was used to measure Gs coupling in cells expressing OA1.0 or Octβ2R; n = 3 wells containing >30 cells each. (G) Left: Normalized change in fluorescence measured in OA1.0-expressing cells in response to the indicated compounds applied at 10 μM (except Ep, which was applied at 100 μM); n = 3 wells containing >300 cells each. Right: Dose-response curves measured in OA1.0-expressing cells in response to OA, tyramine (TA), dopamine (DA), and norepinephrine (NE), with the corresponding EC_50_ values shown; n = 3 wells containing >300 cells each. ACh, acetylcholine; Glu, glutamate; GABA, γ-aminobutyric acid. (H) Left: Exemplar cyclic voltammograms for 100 μM OA, TA, DA, and NE measured using fast-scan cyclic voltammetry (FSCV); the traces were averaged from separate trials. Right: The voltammetric current responses at 0.6 V were measured in accordance with the increasing concentrations of OA, TA, DA, and NE; the inset shows the summary data in response to 100 μM OA, TA, DA, and NE. (I) Schematic illustration depicting the *in vivo* imaging setup using and perfusion to the brain of flies expressing OA1.0 in the mushroom body (MB, 30y-GAL4-driven). (J) Representative *in vivo* fluorescence images (top left), pseudocolor images (top right), traces (bottom left), and summary (bottom right) of the change in OA1.0 fluorescence measured in the MB horizontal lobe in response to application of DA (500 μM), TA (500 μM), OA (500 μM), and Ep (100 μM). In this and subsequent Fig.s, all summary data are presented as the mean ± SEM, superimposed with individual data. *p < 0.05, ***p < 0.001, and n.s., not significant (for F, G, and H, one-way ANOVA with Tukey’s post hoc test; for J, paired or unpaired Student’s t-test). Scale bar = 20 μm.

When expressed in HEK293T cells, OA1.0 trafficked to the plasma membrane and produced a peak change in fluorescence (ΔF/F_0_) of ∼660% in response to 100 μM OA (Fig. 1C). To measure the sensor’s kinetics, we used a rapid perfusion system to locally apply OA followed by the OA receptor antagonist epinastine (Ep), and we measured the change in fluorescence using high-speed line scanning. The data were then fitted to obtain an on-rate (τon) and off-rate (τoff) of approximately 0.02 s and 1.40 s, respectively (Fig. 1D). We also measured the spectral properties of OA1.0 using both one-photon (1P) and two-photon (2P) excitation, which revealed excitation peaks at ∼500 nm and ∼920 nm, respectively, and an emission peak at ∼520 nm (Fig. 1E), similar to those of other commonly used green fluorescent probes. To confirm that OA1.0 does not activate signaling pathways downstream of Octβ2R (thus not affecting cellular physiology), we measured β-arrestin and Gs pathway activation using the Tango assay and the red cAMP sensor RFlamp, respectively. Cells expressing OA1.0 exhibited negligible β-arrestin-dependent signaling compared to cells expressing WT Octβ2R, even at high concentrations of OA (Fig. 1F, left). Moreover, cells expression OA1.0 had significantly lower downstream Gs coupling compared to cells expressing WT Octβ2R (Fig. 1F, right).

With respect to its specificity, we found that the OA1.0 signal induced by OA was abolished by Ep, and the application of several other neurotransmitters did not produce a detectable change in fluorescence (Fig. 1G, left). Next, we measured the response of OA1.0 to various concentrations of OA, as well as the structurally similar transmitters tyramine (TA), dopamine (DA) and norepinephrine (NE). We found that OA1.0 has an ∼40-fold higher affinity for OA (EC_50_ = ∼200 nM) compared to TA (EC_50_ = ∼8000 nM), and showed a negligible response to DA and NE at all tested concentrations (Fig. 1G, right). However, the utilization of the FSCV method for OA detection does not offer such robust specificity, as we observed significant interference from DA and NE in OA detection despite the relatively minor disruption from TA (Fig. 1H).

To evaluate the specificity of OA1.0 *in vivo*, we generated transgenic flies expressing OA1.0 in the MB (30y-GAL4-driven) and then sequentially applied DA, TA, OA and Ep to the fly brain while performing 2P imaging. We found that neither DA nor TA induced an obvious response, while OA elicited a robust response in OA1.0 fluorescence (with a peak ΔF/F_0_ of ∼100%) that was blocked by Ep (Fig. 1I and J). Together, these data demonstrate that OA1.0 can reliably measure the dynamics of OA release with high specificity for OA.

### OA1.0 can report endogenous OA release signals *in vivo*

To further characterize the release of endogenous OA *in vivo*, we used *Drosophila* expressing OA1.0 in the MB (MB247-LexA-driven), which receives projections from several pairs of OANs, including ventral unpaired median a2 (VUMa2) neurons, ventral paired median 3 (VPM3) neurons, VPM4 neurons, VPM5 neurons, and APL neurons[23, 54].To induce the release of endogenous OA in the MB, we applied local electrical stimuli at 30 Hz and observed an incremental increase in fluorescence with an increasing number of stimuli, and this response was eliminated by Ep (Fig. 2A-2D). Moreover, the response was specific to OA, as no detectable response to electrical stimuli was measured in flies lacking TβH in the OANs (Tdc2-GAL4-driven) (Fig. 2C and D). When we applied 50 electrical stimuli at a frequency of 100 Hz, we measured τ_on_ and τ_off_ rates of ∼0.6 s and ∼9.4 s, respectively (Fig. 2E).

**Figure 2.**
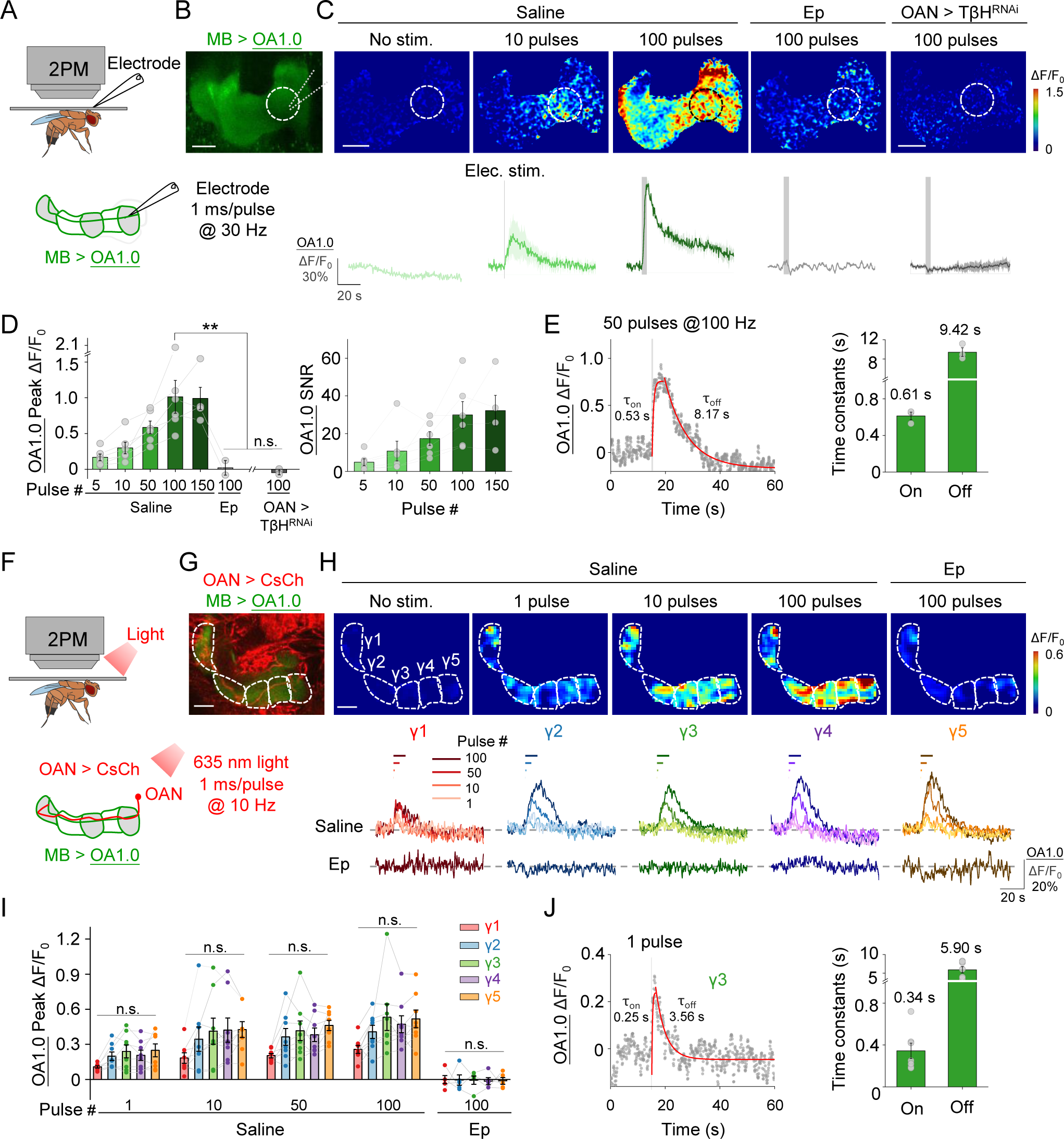
OA1.0 can report the release of OA release *in vivo*. (A) Schematic illustration depicting the experimental setup in which a transgenic fly expressing OA1.0 in the MB (MB247-LexA-driven) is fixed under a two-photon microscope (2PM) and a glass electrode is used to apply electrical stimuli near the MB. (B) Example fluorescence image of OA1.0 expressed in the MB. The dotted circle represents the region of interest (ROI) used for subsequent analysis. (C) Representative pseudocolor images (top) and corresponding traces (bottom) of the change in OA1.0 fluorescence in response to the indicated number of electrical stimuli in a control fly, a control fly treated with 100 μM epinastine (Ep), and an OAN (Tdc2-GAL4-driven) > TβH^RNAi^ fly. (D) Summary of peak ΔF/F_0_ (left) and the signal-to-noise ratio (SNR, right) measured in response to electrical stimuli for the indicated conditions; n = 2-6 flies/group. (E) Left: Time course of ΔF/F_0_ measured in OA1.0-expressing flies in response to 50 electrical stimuli applied at 100 Hz; the rise and decay phases were fitted with a single-exponential function (red traces). Right: Summary of τ_on_ and τ_off_; n = 3 flies/group. (F) Schematic illustration depicting the experimental setup for optogenetic stimulation. (G) Example dual-color fluorescence image of OA1.0 expressed in the MB (green, MB247-LexA-driven) and CsChrimson-mCherry expressed in OANs (red, Tdc2-GAL4-driven); the ΔF/F_0_). The γ1-γ5 compartments of the MB are indicated using dashed lines. (H) Representative pseudocolor images (top) and corresponding traces (bottom) of the change in OA1.0 fluorescence measured in response to the indicated number of optogenetic stimuli applied either in saline or 100 μM Ep. (I) Summary of peak ΔF/F_0_ measured in response to optogenetic stimuli; n = 8 flies/group. (J) Left: Time course of ΔF/F_0_ measured in the γ3 compartment in response to a single laser pulse; the rise and decay phases were fitted with a single-exponential function (red traces). Right: Summary of τ_on_ and τ_off_; n = 7 flies/group. **p < 0.01, and n.s., not significant (for D, paired or unpaired Student’s t-test; for I, one-way ANOVA with Tukey’s post hoc test). Scale bar = 20 μm.

To monitor the release of OA in response to the direct activation of OANs *in vivo*, we optogenetically activated OANs (Tdc2-GAL4-driven) in flies expressing CsChrimson-mCherry while simultaneously imaging OA1.0 expressed in the MB (MB247-LexA-driven) (Fig. 2F, 2G). We found that activating OANs induced a transient increase in OA1.0 fluorescence in the γ1-γ5 compartments of the MB, with the magnitude of the OA1.0 response dependent on the number of light pulses applied; moreover, the peak responses were similar among all five γ compartments (Fig. 2H and I). Importantly, the response for 100 pulses stimulation was blocked in all five compartments by Ep, confirming the sensor’s specificity (Fig. 2H and I). We then measured the kinetics of the response using the γ3 compartment as an example and found that a single pulse of 635-nm laser evoked a measurable increase in OA1.0 fluorescence, with τon and τoff values of ∼0.34 s and ∼5.90 s, respectively (Fig. 2J). Taken together, these results show that OA1.0 can be used *in vivo* to monitor endogenous OA release with high spatiotemporal resolution, high specificity, and high sensitivity.

### OA1.0 can detect physiologically evoked OA release in the MB of living flies

The conflicting findings regarding the role of OA in aversive olfactory learning[19, 28, 29] highlight the need to better understand whether OA release can be activated by odor and/or an aversive stimulus such as electric body shock, which can represent either the CS or the US in this type of learning. To address this question, we expressed OA1.0 in the *Drosophila* MB (MB247-LexA-driven) and found that both odorant application and electric body shock induced a time-locked increase in OA1.0 fluorescence in all five γ compartments, with no difference observed among the various compartments (Fig. 3A-C). In contrast, we found no detectable response to either odorant application or electrical shock in flies in which we knocked down TβH expression in OANs or in flies which OAN activity was suppressed by expressing the inward rectifying potassium channel Kir2.1. As an internal control, direct application of OA still elicited a robust OA1.0 response in both models (Fig. S2).

**Figure 3.**
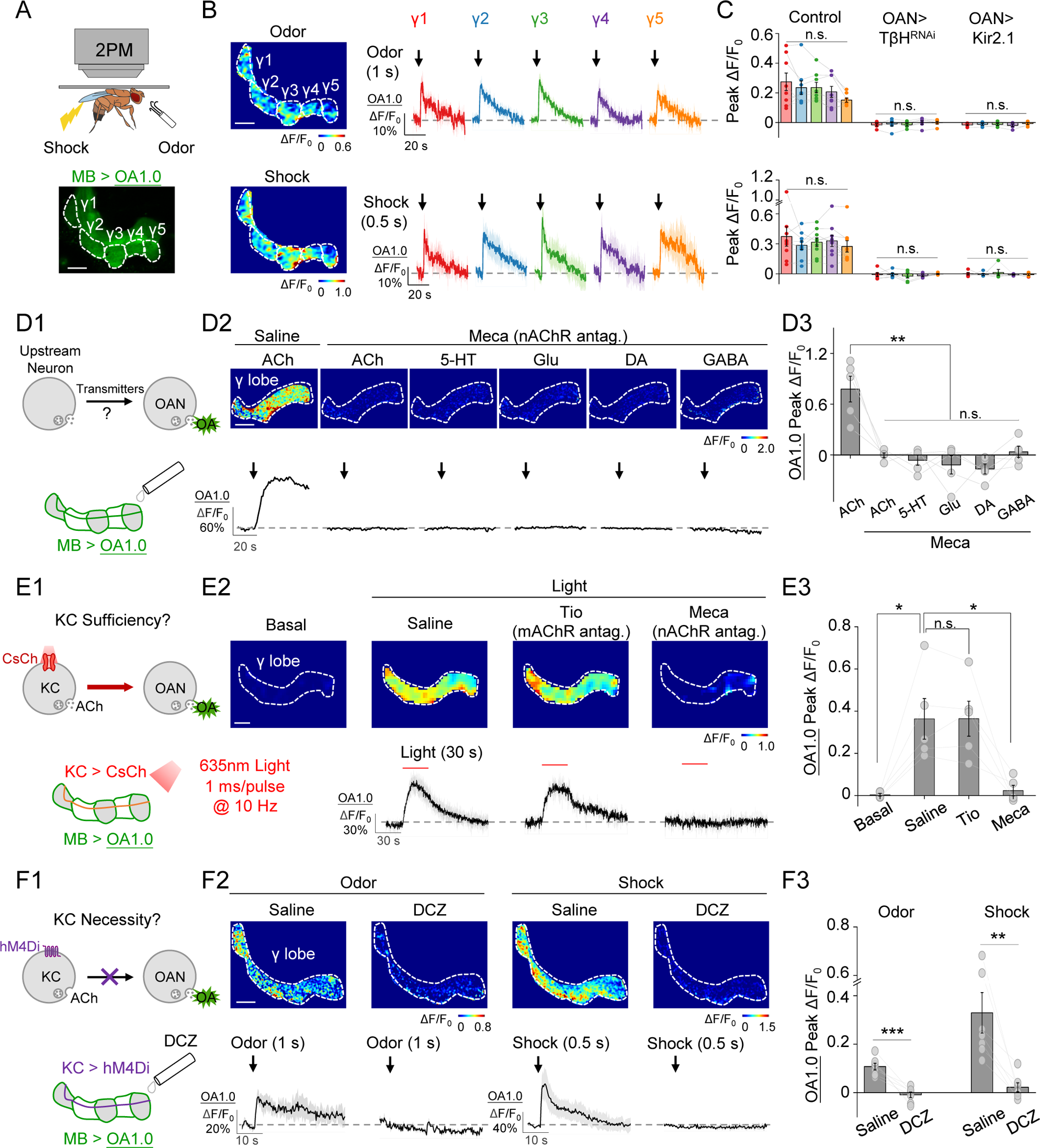
OA1.0 reveals that OA release induced by odor and shock stimuli is activated by ACh released from KCs. (A) Schematic diagram depicting the experimental setup for 2PM with odor and body shock stimulation in flies expressing OA1.0 in the MB (MB247-LexA-driven), with an example fluorescent image of the MB shown below. (B-C) Representative pseudocolor images (B, left), traces (B, right), and summary (C) of the change in OA1.0 fluorescence measured in response to odorant application (top) and body shock (bottom) in OA1.0-expressing flies (n = 8-9) and OA1.0-expressing flies co-expressing TβH^RNAi^ (n = 6) or Kir2.1 (n = 5) in OANs (Tdc2-GAL4-driven). (D) Schematic diagram (D1) depicting the strategy used to apply compounds to the brain of flies expressing OA1.0 in the MB (MB247-LexA-driven). Also shown are representative pseudocolor images (D2, top), traces (D2, bottom), and summary (D3) of the change in OA1.0 fluorescence in response to the indicated compounds (1 mM each) applied in the absence or presence of the nAChR antagonist Meca (100 μM); n = 5 flies/group. (E) Schematic diagram (E1) depicting the strategy in which CsChrimson expressed in KCs (R13F02-GAL4-driven) was activated using optogenetic stimulation, and OA1.0 fluorescence was measured in the MB (MB247-LexA-driven). Also shown are representative pseudocolor images (E2, top), traces (E2, bottom), and summary (E3) of the change in OA1.0 fluorescence in response to optogenetic stimulation in saline, the muscarinic ACh receptor antagonist Tio (100 μM), and Meca (100 μM); n = 5 flies/group. (F) Schematic diagram (F1) depicting the strategy in which hM4Di expressed in KCs (30y-GAL4-driven) was silenced by applying 30 nM deschloroclozapine (DCZ), and OA1.0 fluorescence was measured in the MB. Also shown are representative pseudocolor images (F2, top), traces (F2, bottom), and summary (F3) of the change in OA1.0 fluorescence in response to odor or electrical body shock in the absence or presence of 30 nM DCZ; n = 7 flies/group. *p < 0.05, **p < 0.01, ***p < 0.001, and n.s., not significant (for C, one-way ANOVA with Tukey’s post hoc test; for D3-F3, paired Student’s t-test). Scale bar = 20 μm.

### OA1.0 reveals that KC activity is both necessary and sufficient for OA release in the *Drosophila* MB

Next, to examine the mechanism underlying OA release in the MB, we attempted to identify the neurons and pathways that regulate OAN activity. Although previous connectomic analyses showed that KCs, the principal neurons in the MB, are the primary cells upstream of OANs (Fig. S3)[55, 56], the functional inputs that drive OA release are currently unknown. Given that KCs release the excitatory neurotransmitter acetylcholine (ACh)[57], we perfused ACh onto the γ lobe of the MB and observed an increase in OA1.0 fluorescence that was prevented by the nicotinic ACh receptor (nAChR) antagonist mecamylamine (Meca). Moreover, we found no increase in OA1.0 fluorescence when other neurotransmitters such as 5-hydroxytryptamine (5-HT), glutamate (Glu), DA and γ-aminobutyric acid (GABA) were applied in the presence of Meca (Fig. 3D).

Because perfusion of exogenous ACh lacks cell-type specificity, we used optogenetics to determine whether selectively activating KCs (R13F02-GAL4-driven) is sufficient to induce OA release in the MB. Consistent with our perfusion experiments, we found that optogenetically activating KCs caused an increase in OA1.0 fluorescence that was blocked by Meca but not the muscarinic ACh receptor antagonist tiotropium (Fig. 3E). Moreover, there is no obvious light-induced OA release in transgenic flies with UAS-CsChrimosn but without KC-GAL4 (R13F02-GAL4) (Fig. S4A), ruling out the unspecific effect due to the leaky expression of channelrhodopsin[58]. Together, these results suggest that ACh release from KCs serves as the excitatory signal that drives OA release via nAChRs in the γ lobe of the MB.

To determine whether KCs are required for activating OANs in the MB, we generated transgenic flies expressing both OA1.0 and the inhibitory DREADD (designer receptors exclusively activated by designer drugs) hM4Di[59–61], and found that both odor- and shock-induced OA1.0 signals were abolished when KCs activity was suppressed by the hM4Di agonist deschloroclozapine (DCZ)[62] (Fig. 3F). Meanwhile, the DCZ application showed no significant effect on stimuli-induced OA signals in flies without hM4Di (Fig. S4B). Thus, KC activity is both necessary and sufficient for OA release from OANs in the MB.

### OA regulates aversive learning behavior and related synaptic plasticity

To examine the biological significance of OA release triggered by odorant application and body shock, we measured aversive learning and the coincident time window in flies lacking either OA synthesis or OAN activity. We found that both TβH mutant flies and OAN-silenced flies expressing Kir2.1 had significantly reduced learning performance compared to WT flies (Fig. 4A and B). Moreover, unlike flies lacking neuronal tryptophan hydroxylase (Trhn), the rate-limiting enzyme in 5-HT biosynthesis, which have a significantly shortened coincident time window compared to control flies, the coincident time window was unchanged in TβH mutants (Fig. S5). These results suggest that OA plays a key and specific role in aversive learning ability in *Drosophila*.

**Figure 4.**
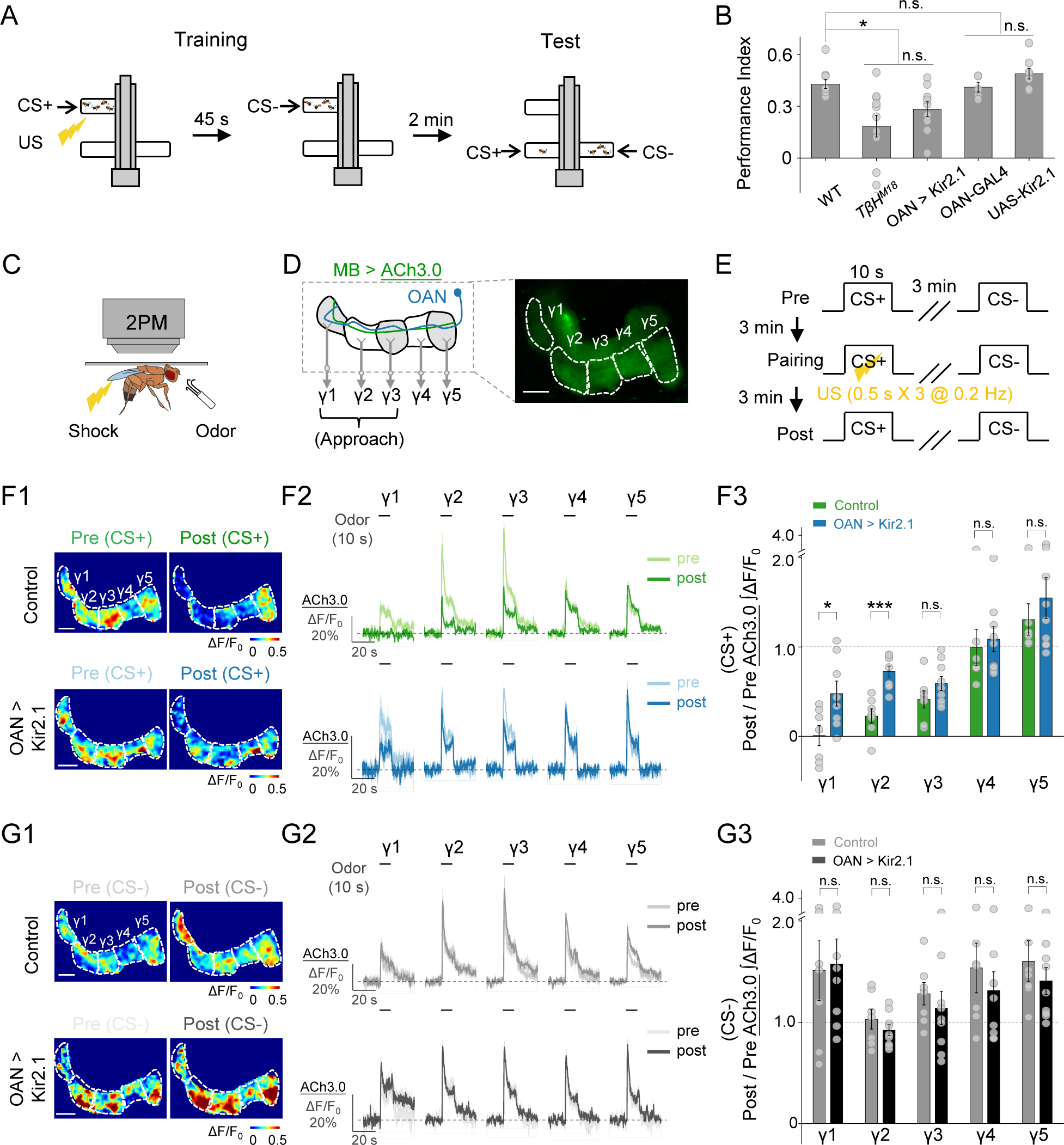
OA plays an essential role in aversive learning and synaptic plasticity in KCs in the MB. (A) Schematic diagram depicting the T-maze protocol for measuring aversive learning in *Drosophila*. (B) Summary of the performance index measured in WT flies and the indicated transgenic flies. OAN-GAL4 and UAS-Kir2.1 served as control groups; n=5-10 for each group. (C-E) Schematic diagram (C) depicting the *in vivo* 2PM imaging setup, a representative fluorescence image (D), and the experimental protocol (E) in which odor-induced changes in ACh3.0 fluorescence (MB247-LexA-driven) in the γ1-γ5 compartments were measured before (pre), during, and after (post) pairing. (F-G) Representative pseudocolor images (F1, G1) and average traces (F2, G2) of odor-evoked ACh3.0 responses measured in the γ1-γ5 compartments before and after pairing in response to the CS+ odorant (F) and CS- odorant (G) in control flies (top) and OAN-silenced (OAN > Kir2.1) flies (bottom). F3 and G3: Summary of the change in odor-evoked ACh release (post/pre responses) after pairing in response to the CS+ odorant (F3) and CS- odorant (G3) in control flies and OAN > Kir2.1 flies; n = 6-9 flies/group. *p < 0.05, ***p < 0.001, and n.s., not significant (unpaired Student’s t-test). Scale bar= 20 μm.

Given that synaptic plasticity is fundamental to the neuronal basis of learning, the regulation of synaptic plasticity by OAN activity after odor-shock pairing is a potential mechanism underlying the observed aversive learning results. Previous electrophysiological recordings or Ca^2+^ imaging studies in the mushroom body output neuron (MBON) innervating the γ1 compartment (MBON-γ1pedc) suggested that pairing an odorant with dopaminergic reinforcement induces synaptic depression between KCs and the MBON[63–65]. This synaptic depression is correlated with decrease ACh release from KCs[66, 67]. Thus, we used the GRAB_ACh3.0_ sensor (ACh3.0)[45] to monitor the ACh release in the γ lobe of the MB (MB247-LexA-driven) (Fig. 4C-4E). By comparing the odor-evoked ACh release measured before and after odor-shock pairing in control flies, we observed significant synaptic depression in the γ1, γ2 and γ3 compartments (Fig. S6), the three compartments known to transmit information to MBONs associated with approach behavior[68]. We then examined ACh release following odor-shock pairing in flies expressing Kir2.1 in the OANs. Our results revealed significantly less synaptic depression (i.e., reduced depression of ACh release) in the CS+ response, specifically in the γ1 and γ2 compartments compared to control flies (Fig. 4F), indicating impaired synaptic plasticity during learning in OAN-silenced flies. In contrast, we found no significant difference in the change in ACh release in response to CS- (a separate odorant that was not paired to the electric body shock) between OAN-silenced flies and control flies in any γ compartments (Fig. 4G). Taken together, these results suggest that OA plays an essential role in modulating the change in synaptic plasticity induced by odor-shock pairing, thereby amplifying the aversive learning behavior.

### OA regulates aversive learning by modulating US processing via Octβ1R expressed on dopaminergic neurons

Synchronization between the CS and the US is required for aversive learning; specifically, information regarding the CS is conveyed by projection neurons to the calyx of the MB for processing by KCs, while information regarding the US is conveyed by dopaminergic neurons (DANs) to the MB lobes for subsequent processing[69]. We therefore examined the effect of OA on CS and/or US processing in regulating aversive learning. For this experiment, we expressed the calcium sensor GCaMP6s in KCs (MB247-LexA-driven) to measure calcium signals in the calyx, thus providing information regarding the dynamics of CS processing (Fig. 5A1). In separate experiments, we expressed the GRAB_DA2m_ (DA2m) sensor[47] in the MB (R13F02-LexA-driven) to measure DA release in the γ lobe, thus capturing the dynamics of US processing (Fig. 5B1). In both cases, we used both control flies and OAN-silenced flies to specifically examine the role of OA in aversive learning. We found that the calcium signals measured in the calyx in response to odorant application were similar between OAN-silenced flies and control flies (Fig. 5A2 and A4); in contrast, shock-induced DA release in the γ lobe was significantly lower in OAN-silenced flies (Fig. 5B3 and B4). Notably, we found that the shock stimuli induced small calcium signals in the KCs of the calyx, while odor stimuli induced small DA transients in the γ lobe; moreover, no significant differences were observed in these responses between OAN-silenced flies and the corresponding control flies (Fig. 5A3, A4, B2 and B4). Together, these findings suggest that OAN activity modulates US processing, but not CS processing, during aversive learning.

To eliminate potential developmental influences on our observations regarding the effect of OA on DA release in response to the US, we applied the OA receptor antagonist Ep to the fly’s brain and found that the same individual fly exhibited a significant reduction in shock-induced DA release along the γ lobe compared before and after the Ep treatment (Fig. 5C, left and middle). Previous studies showed that short-term aversive memory formation requires OA signaling via Octβ1R[25]; we therefore specifically knocked down Octβ1R expression in DANs (TH-GAL4-driven) using RNAi (Fig. 5C, right) to examine whether OA directly affects DA release and found a significant decrease in DA release compared to controls (Fig. 5C, left and right). Based on these results, we then examined whether knocking down Octβ1R expression in DANs affects synaptic plasticity and/or learning. Similar to our results obtained with OAN-silenced flies (see Fig. 4), we found significant differences in the degree of KC synaptic depression in response to CS+ in both the γ1 and γ2 compartments of Octβ1R-knockdown flies compared to control flies. In contrast, we found no significant differences in the γ3, γ4, or γ5 compartments in response to CS+, or in any γ compartment in response to CS- (Fig. 6A-6E). Moreover, both Octβ1R-knockout flies and Octβ1R-knockdown flies displayed significantly impaired learning compared to control flies (Fig. 6F). These results support a model in which OA boosts aversive learning via Octβ1R in DANs, which enhances the punitive US signals to modulate synaptic plasticity in KCs (Fig. 6G).

**Figure 5.**
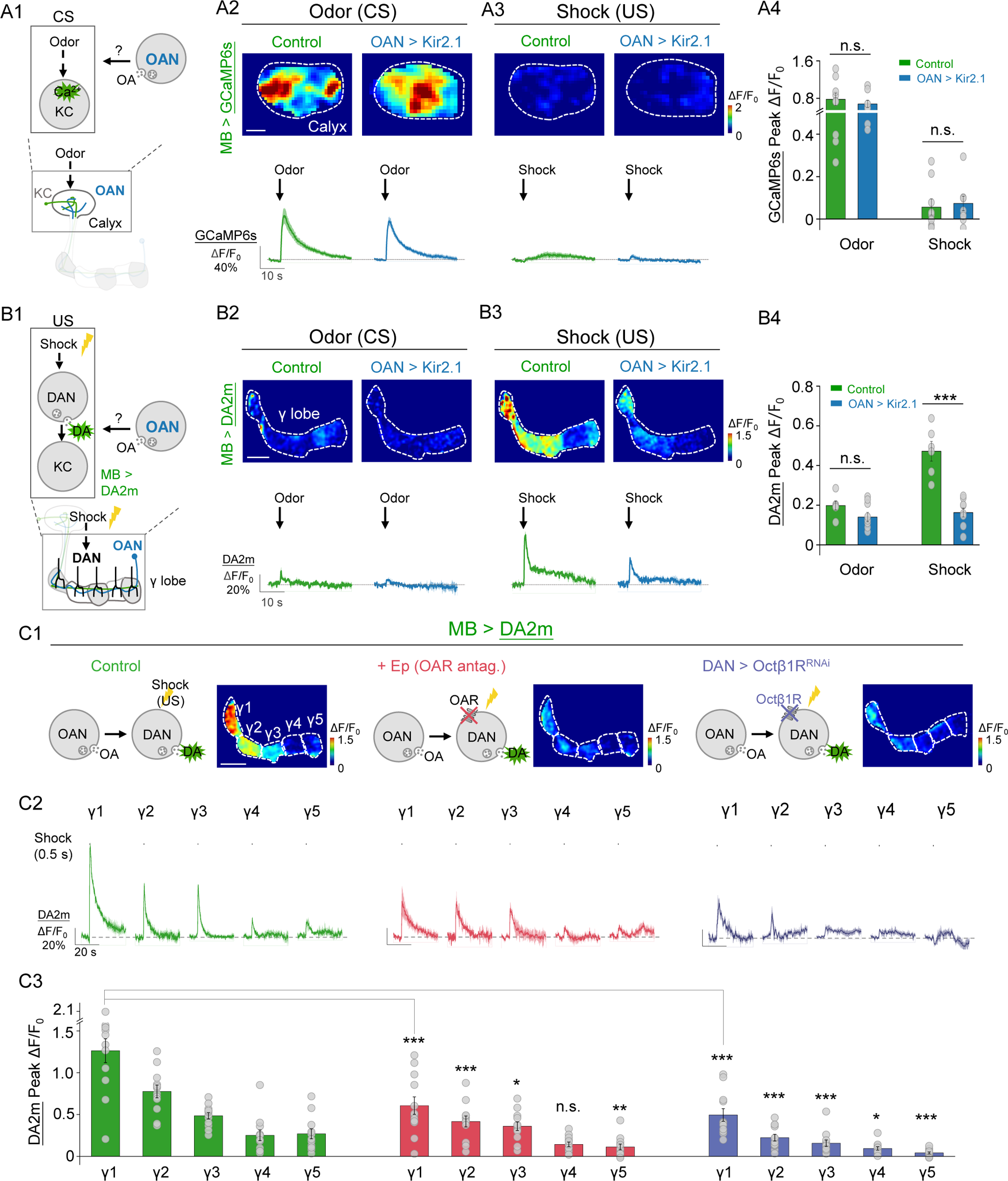
OA is required for driving DA release in response to aversive stimuli. (A) Schematic diagram (A1) showing the strategy for measuring intracellular calcium signals in the MB (MB247-LexA-driven) by expressing GCaMP6s in either control flies or OAN > Kir2.1 flies, in response to the conditioned stimulus (CS) or unconditioned stimulus (US). Also shown are representative pseudocolor images (A2-A3, top), traces (A2-A3, bottom), and summary (A4) of calcium signals measured in the calyx in response to odor (A2) or electrical body shock (A3); n = 9 flies/group. (B) Schematic diagram (B1) showing the strategy for measuring dopamine (DA) signals in the MB (R13F02-LexA-driven) by expressing the DA2m sensor in either control flies or OAN > Kir2.1 flies, in response to the CS or US. Also shown are representative pseudocolor images (B2-B3, top), traces (B2-B3, bottom), and summary (B4) of DA release measured in the γ lobe in response to in response to odor (B2) or electrical body shock (B3); n = 6-9 flies/group. (C) C1: Schematic diagrams (C1) showing DA2m imaging in flies and representative pseudocolor images whose brain was bathed in saline (left) or saline containing 100 μM Ep (middle), or DAN > Octβ1R^RNAi^ (TH-GAL4-driven) flies (right) in response to body shock stimuli. Also shown are representative traces (C2) and the summary (C3) of DA release measured in the γ1-γ5 compartments; n = 12 flies/group. *p < 0.05, **p < 0.01, ***p < 0.001, and n.s., not significant (unpaired Student’s t-test). Scale bar= 20 μm.

**Figure 6.**
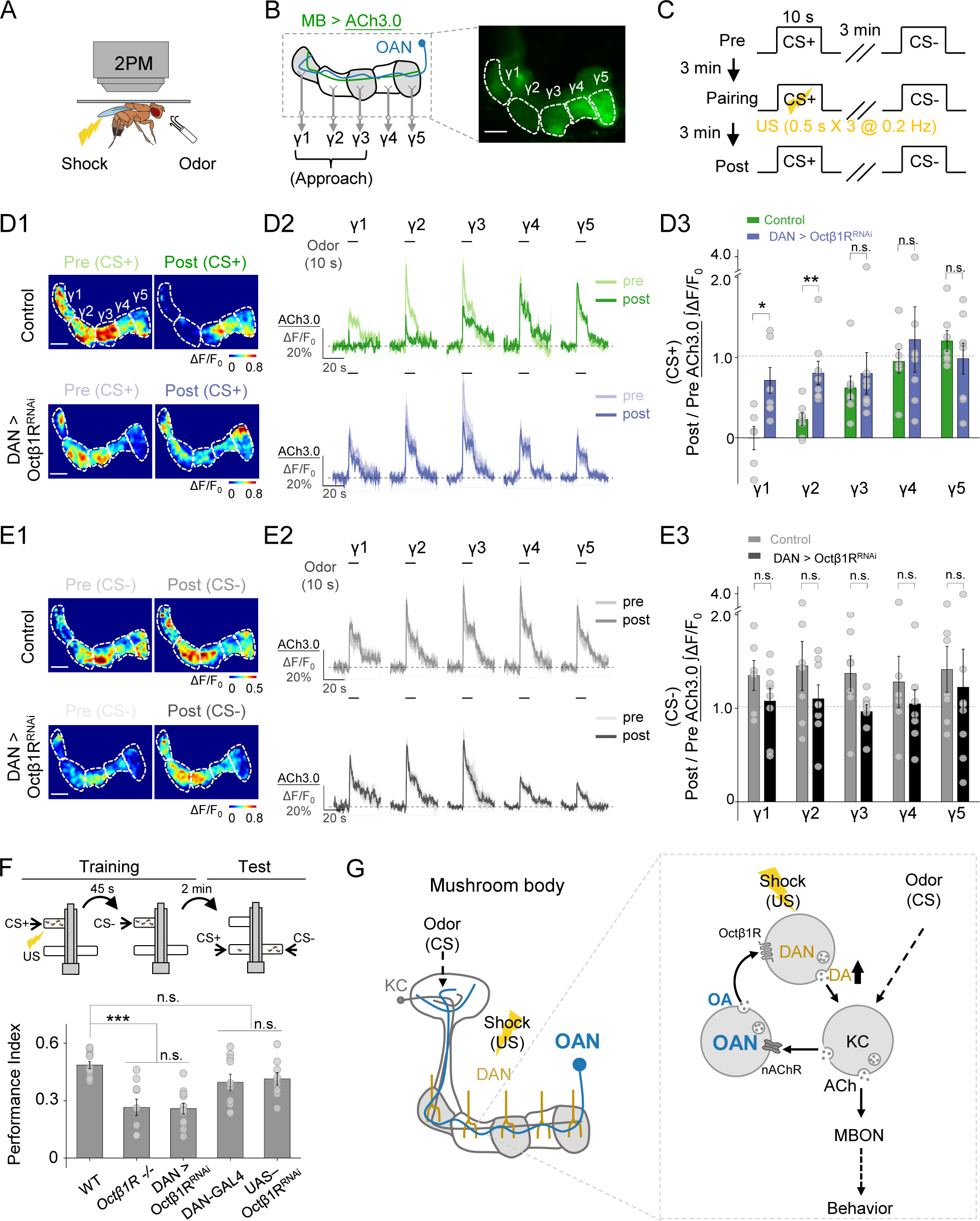
OA acts on DANs via the Octβ1R receptor to modulate aversive learning. (A-C) Schematic diagram (A) depicting the *in vivo* 2PM imaging setup, a representative fluorescence image (B), and the experimental protocol (C) in which odor-induced changes in ACh3.0 (MB247-LexA-driven) fluorescence were measured in the γ1-γ5 compartments before, during, and after pairing. (D-E) Representative pseudocolor images (D1, E1), average traces (D2, E2), and summary (D3, E3) of odor-evoked ACh3.0 responses measured in the γ1-γ5 compartments in response to the CS+ odorant (D) and CS- odorant (E) in the indicated groups; n = 6-8 flies/group. (F) Schematic diagram depicting the T-maze protocol (top) and summary of the performance index (bottom) measured in the indicated groups; n = 9-12 for each group. (G) Model depicting the proposed mechanism for how OA acts on DANs in the MB to modulate aversive learning. MBON, mushroom body output neuron. *p < 0.05, **p < 0.01, ***p < 0.001, and n.s., not significant (unpaired Student’s t-test). Scale bar = 20 μm.

## DISCUSSION

Here, we developed a new genetically encoded fluorescent sensor called GRAB_OA1.0_ to detect OA release with high selectivity, sensitivity, and spatiotemporal resolution both *in vitro* and *in vivo*. We then used this tool to perform the first detailed study of the spatial and temporal dynamics of OA during aversive learning in *Drosophila*. We found that ACh released from KCs activates OANs, triggering OA release via nAChRs. Notably, we also observed that ACh released from KCs is required for OA release in response to both the CS and the US during aversive learning. Furthermore, by integrating other genetically encoded fluorescent sensors (namely, GRAB_DA2m_ and GRAB_ACh3.0_ to monitor DA and ACh, respectively), we discovered that OA increases shock-induced DA release via Octβ1R, which in turn regulates the corresponding changes in synaptic plasticity in the MB, ultimately facilitating aversive learning.

### Advantages of OA1.0 over other methods for measuring OA

Compared to other methods used to measure OA, OA1.0 offers several advantages. First, OA1.0 exhibits high specificity for OA over most neurotransmitters such as TA, DA and NE. This is particularly important for detecting OA in the presence of other structurally similar molecules, as electrochemical tools like FSCV cannot distinguish between OA and other chemicals, as shown here (Fig. 1H) and in previous studies[39–41]. Second, OA1.0 offers sub-second kinetics and is genetically encoded, allowing for the non-invasive monitoring of octopaminergic activity *in vivo* with a high recording rate. In contrast, microdialysis has relatively low temporal resolution and requires the placement of a relatively large probe, making it unsuitable for use in small model organisms such as *Drosophila*. Capitalizing on these advantages, we used OA1.0 to monitor OA release *in vivo* in response to a variety of stimuli, gaining new insights into the functional role of OA.

Importantly, OA1.0 can also be expressed in other animal models, including mammals, opening up new opportunities to monitor OA dynamics in a wide range of species. In mammals, OA is classified as a trace amine and exerts its activity through trace amine-associated receptors (TAARs). TAAR1, in particular, has been implicated as a key regulator of monoaminergic and glutamatergic signaling in brain regions relevant to schizophrenia, as demonstrated in knockout and overexpression models in rodents[70, 71]. However, studying TAAR1 is challenging due to the presence of various endogenous ligands, including the trace amines β-phenylethylamine (PEA), TA, and OA, as well as the monoamine neurotransmitters DA, 5-HT, and NE[72]. Thus, the development of robust tools like OA1.0 that selectively monitor a given trace amine will advance our understanding of specific TAAR-mediated biological effects. Additionally, this strategy can be employed to develop sensors for detecting other key trace amines, providing valuable information regarding these chemicals’ dynamics under both physiological and pathological conditions.

### OA plays a key role in associative learning

OA was initially believed to play a role only in appetitive learning, but not in aversive learning, in invertebrates such as *Drosophila*, honeybees, and crickets[19, 28, 73, 74]. However, several studies suggest that OA may indeed be involved in aversive learning, albeit without completely understanding the underlying mechanisms and spatiotemporal dynamics[23, 25, 29]. Schwaerzel et al. first showed that OA has the selective role in *Drosophila*, reporting that TβH mutants had impaired appetitive learning but normal aversive learning[19]. However, it is important to note that the TβH mutants used by Schwaerzel et al. were a mixture of homozygous and hemizygous TβH^M18^ flies regardless of sex, as <u>the localization of TβH was to the X chromosome</u> <u>and</u> the homozygous TβH^M18^ females were sterile. Subsequently, Iliadi et al. found that both homozygous TβH^M18^ males and females performed impaired aversive conditioning compared to WT flies and heterozygous TβH^M18^ females[29]. Drawing on these previous reports, we used homozygous TβH^M18^ males and females and obtained results similar to Iliadi et al., supporting the notion that OA is required for aversive learning in *Drosophila*.

Moreover, we found that OA release in the γ lobe of the MB plays a crucial role in facilitating the release of DA via Octβ1R, which is selectively coupled to increase intracellular cyclic AMP levels by OA[75], in response to shock stimuli. This increased release of DA drives a change in synaptic plasticity between KCs and the MBON and promotes aversive learning[63, 65, 76–80]. The finding aligns with prior studies showing that DANs are downstream of OANs in reward-based learning[20, 21, 81], suggesting a conserved role for OA in mediating the DANs’ ability to perceive US signals in both positive and negative learning scenarios. It is noteworthy that our study utilized a DA sensor[47] to specifically detect the release of DA itself, providing a more direct assessment of its potential effects on downstream neurons, rather than measuring DAN activity[20, 21]. In addition to confirming the involvement of OA in aversive learning, our study also provides novel insights into the underlying input and output circuitry through which OA operates (see Fig. 6G), which potentially indicates that the CS and the US are not entirely independent events within the learning context, but rather, one might have an impact on the other.

Nevertheless, further studies are needed to obtain a more comprehensive understanding of the mechanisms through which OA contributes to associative learning. Notably, previous studies found that Octβ1R, expressed in KCs, is involved in aversive learning[25], which operates as a parallel circuit along with the well-known DA-dDA1 (MB-γ)-MBON pathways[82]. Additionally, in the context of appetitive learning, the α1-like OA receptor OAMB has been shown to play a role in engaging octopaminergic signaling in KCs[22]. These intriguing findings suggest that OA may exert a direct effect on KCs to affect associative learning. Thus, further research is needed in order to unravel the complex interactions and mechanisms by which OA modulates associative learning.

### Neuromodulators interact in associative learning

As the primary center of associative memory in *Drosophila*, the MB uses ACh as the predominant excitatory neurotransmitter released from KCs[57]. However, the MB also receives converging inputs from other neuromodulators such as OA, DA, 5-HT, and GABA. The interactions between these neuromodulator systems, as well as with ACh, are essential for controlling the brain’s states and neuronal computations[55]. Here, we show that odor- or shock-evoked release of OA requires ACh release from KCs, and in turn, increases DA release, thereby forming a positive feedback loop that is required for learning. Recent research has shown that normal DAN synaptic release during learning requires KC input to DAN[83]. In addition, KCs have been shown to activate GABAergic APL neurons[84] and serotoninergic dorsal paired medial (DPM) neurons[67], both of which provide negative feedback to KCs. GABA release from APL neurons is believed to contribute to odor-specific memory through sparse coding[85], while 5-HT release from DPM neurons regulates the coincidence time window of associative learning[67]. Thus, as the predominant neuron type in the MB, KCs not only associate CS and US signals but also regulate a variety of neuromodulators to form local feedback loops. These local reentrant loops allow for moment-by-moment updates of both external (i.e., environmental) and internal information, allowing for the appropriate reconfiguration of the flow of information between KCs and MBONs, thus providing behavioral flexibility and the appropriate responses to change the internal and external states of the organism[86].

The interplay between neuromodulators is both complex and essential for shaping the activity of synaptic circuit elements to drive cognitive processes in both invertebrates and mammals. In this respect, our study provides new insights by highlighting the conserved interaction between OA and DA in invertebrates, offering a valuable framework for understanding the complex interplay between DA and other neurotransmitters in associative learning processes. Additionally, a recent study in mammals showed that continuous interactions and updating between ACh and DA signaling in the nucleus accumbens are critical for regulating the striatal output that underlies the acquisition of Pavlovian learning of reward-predicting cues[87, 88]. Given the similarities between OA-DA interaction in invertebrates and the ACh-DA interaction in mammals, it is reasonable to speculate that such interactions are a fundamental feature of the central nervous system. The discovery that such conserved interactions exist between distinct neuromodulator systems provides valuable new insights into the mechanisms that underlie cognitive processes and may have important implications with respect to developing new therapies for cognitive disorders.

## ACKNOWLEDGMENTS

We thank Yi Rao for providing access to the two-photon microscope. We thank Yi Zhong, Lianzhang Wang and Bohan Zhao for helping with T-maze assay. We thank Jun Chu for providing RFlamp sensor. We thank the imaging core facility of State Key Laboratory of Membrane Biology at Peking University (Ye Liang). We thank the Core Facility of Drosophila Resource and Technology of CAS Center for Excellence in Molecular Cell Science (Wei Wu). We thank the National Center for Protein Sciences at Peking University for support and assistance with the Opera Phenix high content screening system. We thank Seth Tomchik, Stephen Zhang, Andrew Lin, Yan Li, Emmanuel Perisse, and Woo Jae Kim for valuable feedback regarding the manuscript. Finally, we thank Yulin Zhao, Hui Dong, Bin Luo and other Li lab members for helpful suggestions and comments on the manuscript.

This work was supported by grants to Yulong Li from the National Key R&D Program of China (2019YFE011781), the National Natural Science Foundation of China (31925017 and 31871087), the NIH BRAIN Initiative (1U01NS113358 and 1U01NS120824), the Feng Foundation of Biomedical Research, the Clement and Xinxin Foundation, the Peking-Tsinghua Center for Life Sciences, the State Key Laboratory of Membrane Biology at Peking University School of Life Sciences, and the New Cornerstone Science Foundation through the New Cornerstone Investigator Program and the XPLORER PRIZE.

## AUTHOR CONTRIBUTIONS

Y.L. supervised the project. M.L. performed all imaging and behavioral experiments (except as otherwise noted). R.Z. and M.L. analyzed EM data. R.C. and H.W. performed the experiments related to sensor development, optimization and characterization in cultured cells. Y.X. performed FSCV experiments. Y.L. and M.L. wrote the manuscript with input from all other authors.

## KEY RESOURCES TABLE

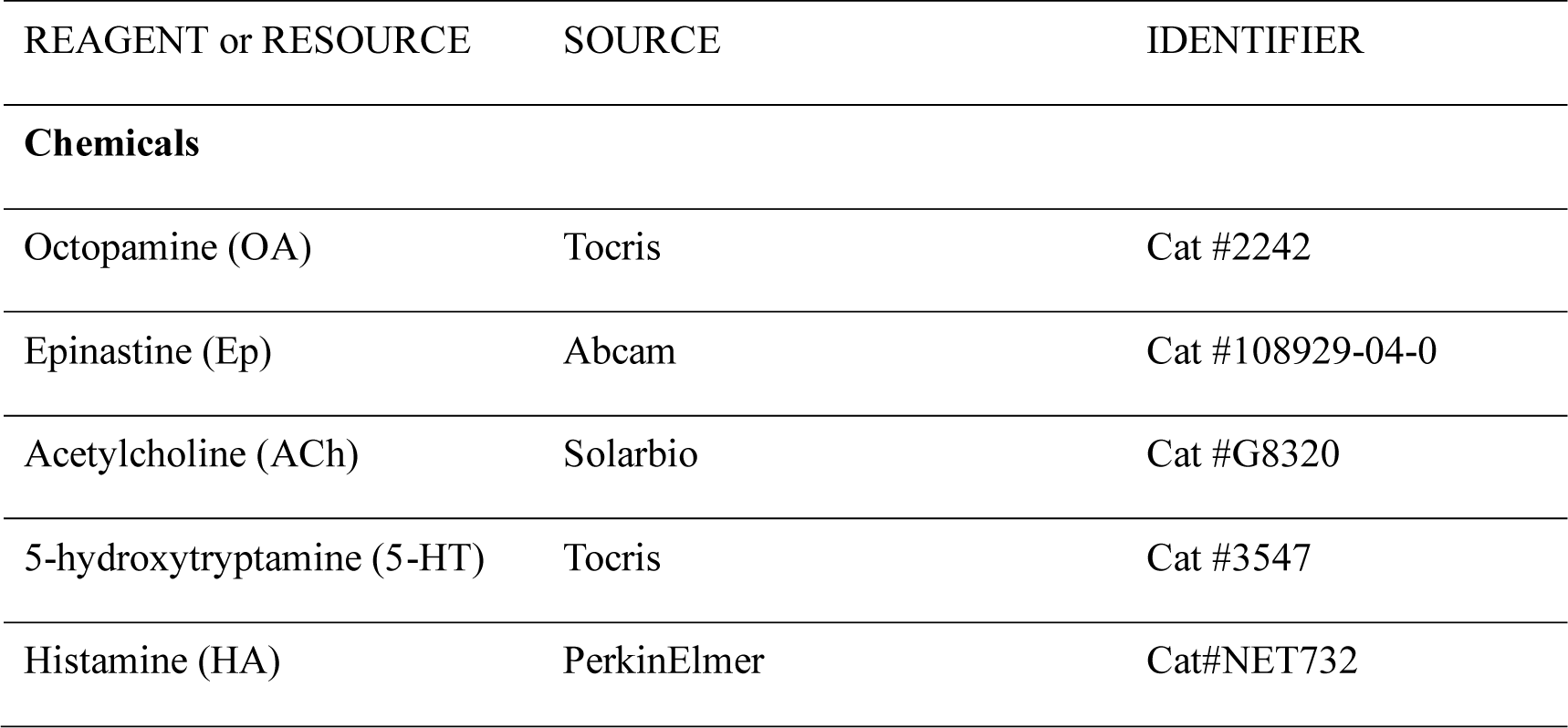
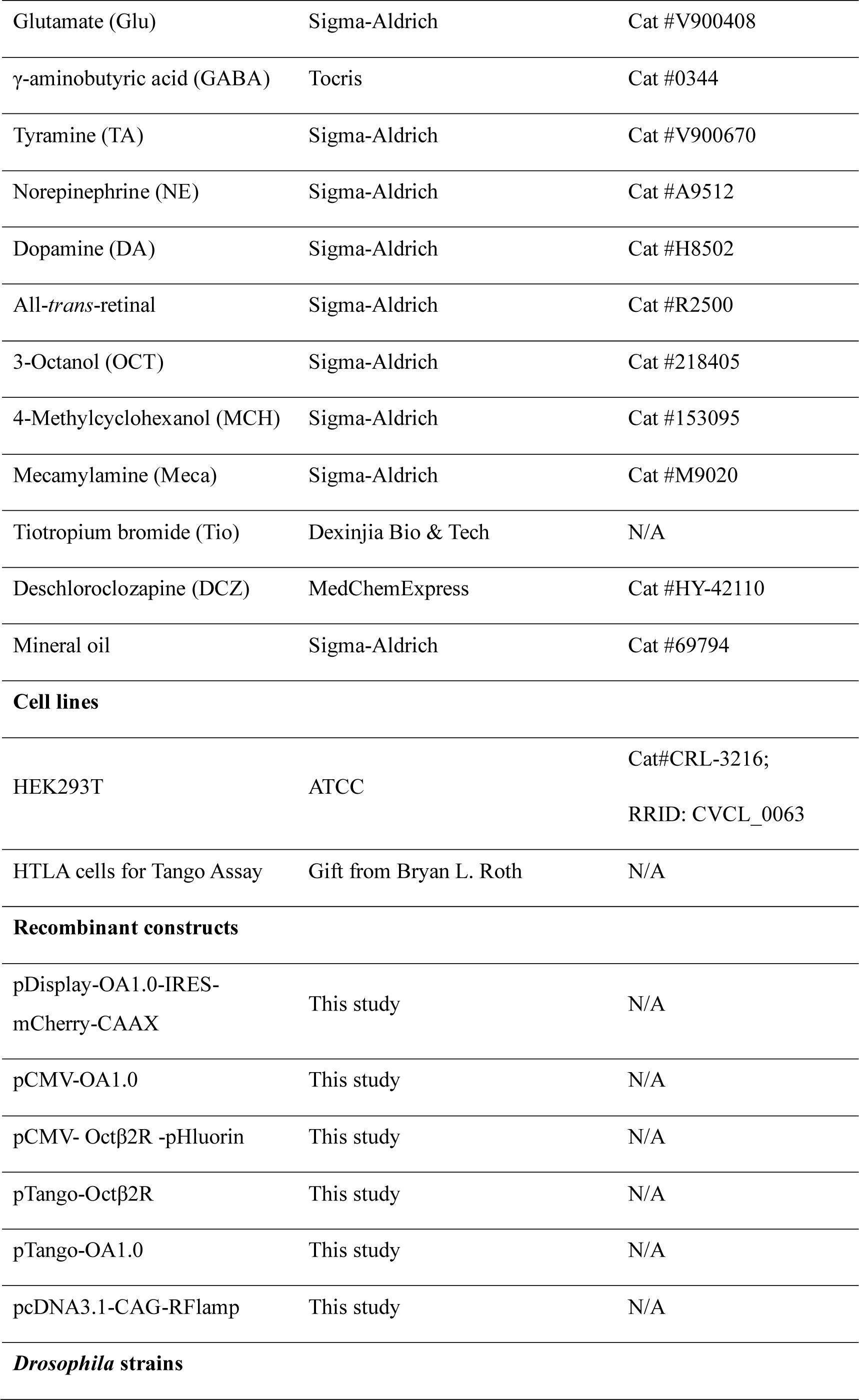
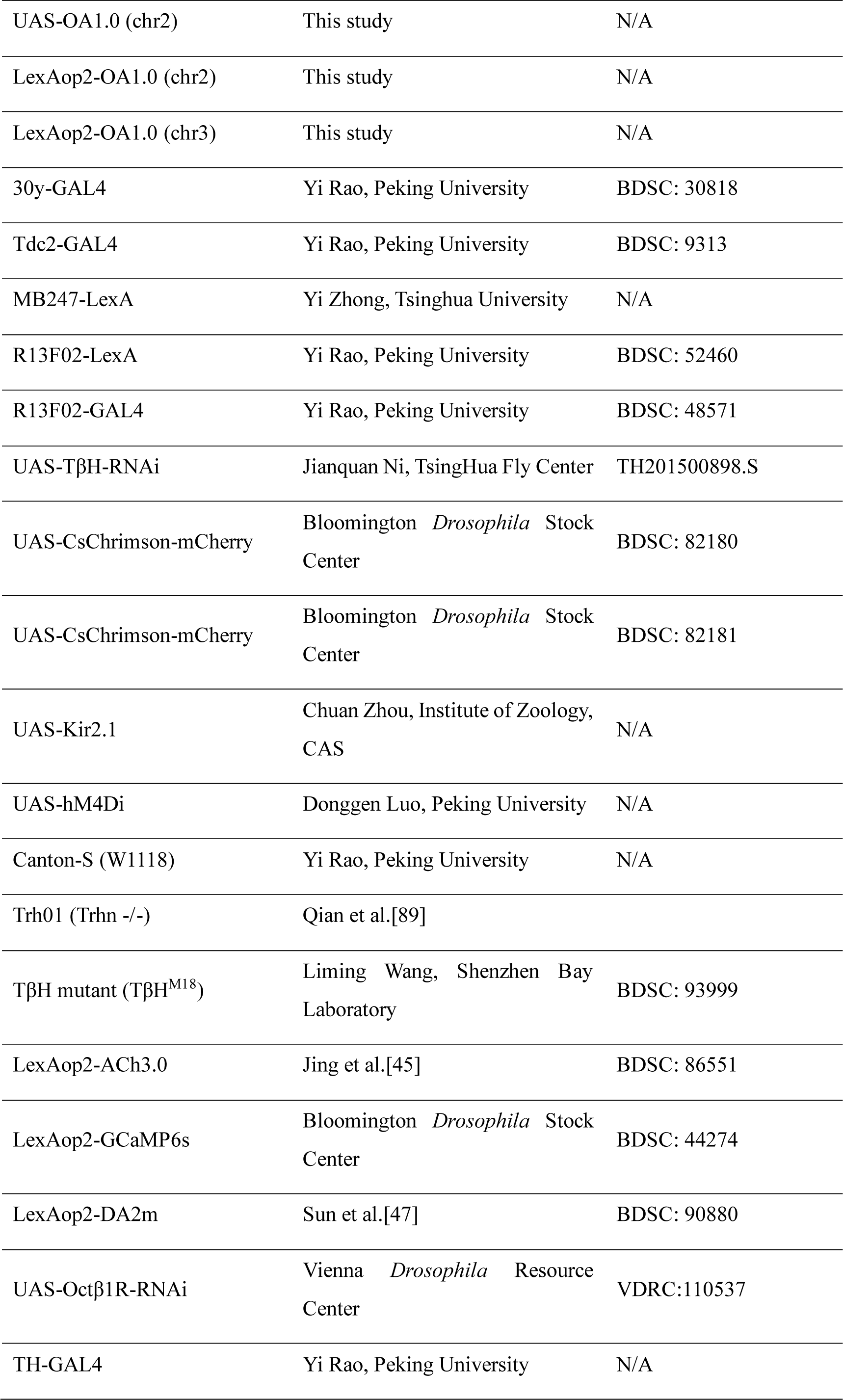
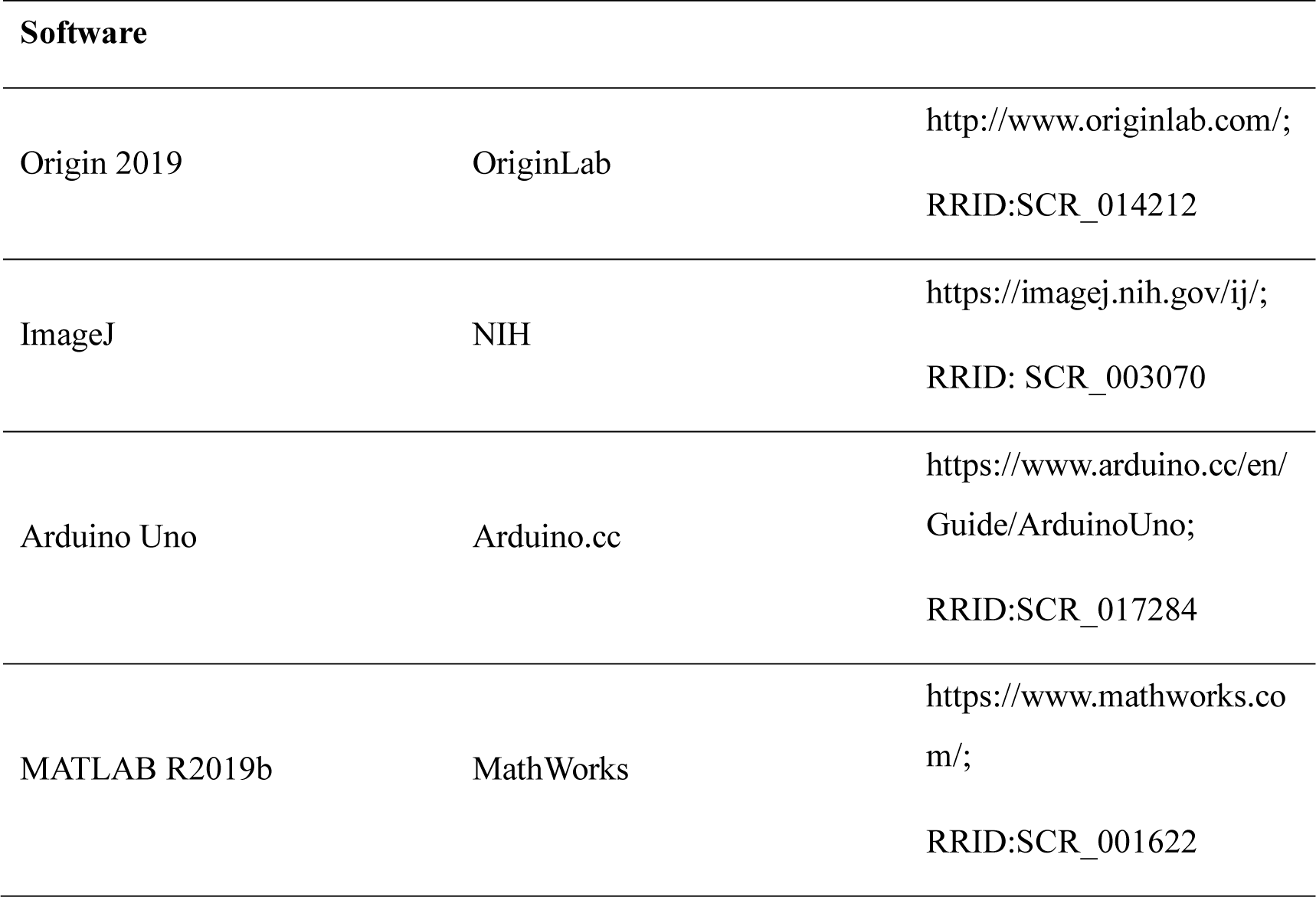

## EXPERIMENTAL MODEL AND SUBJECT DETAILS

### Cell lines

HEK293T cells were acquired from ATCC and verified by microscopic examination of their morphology and growth curve. The cells were cultured in DMEM (Biological Industries) supplemented with 10% (v/v) fetal bovine serum (FBS, Gibco) and 1% penicillin-streptomycin (Gibco) at 37°C in 5% CO_2_.

### Flies

In this study, we generated UAS-OA1.0 (attp40), LexAop2-OA1.0 (attp40) and LexAop2-OA1.0 (vk00005) using Gibson assembly to integrate the coding sequence of OA1.0 into the pJFRC28[90] or modified pJFRC28 vector. The resulting vectors were then injected into *Drosophila* embryos and integrated into attp40 or vk00005 via phiC31 by the Core Facility of Drosophila Resource and Technology, Shanghai Institute of Biochemistry and Cell Biology, Chinese Academy of Sciences.

*Drosophila* were raised at 25°C in 50% humidity and a 12-hour light/dark cycle on a diet of corn meal. For optogenetics experiments, the flies were fed on corn meal containing 400 μM all-trans-retinal immediately after eclosion and kept in total darkness for 8-24 hours prior to imaging experiments.

Detailed fly genotypes used by figures

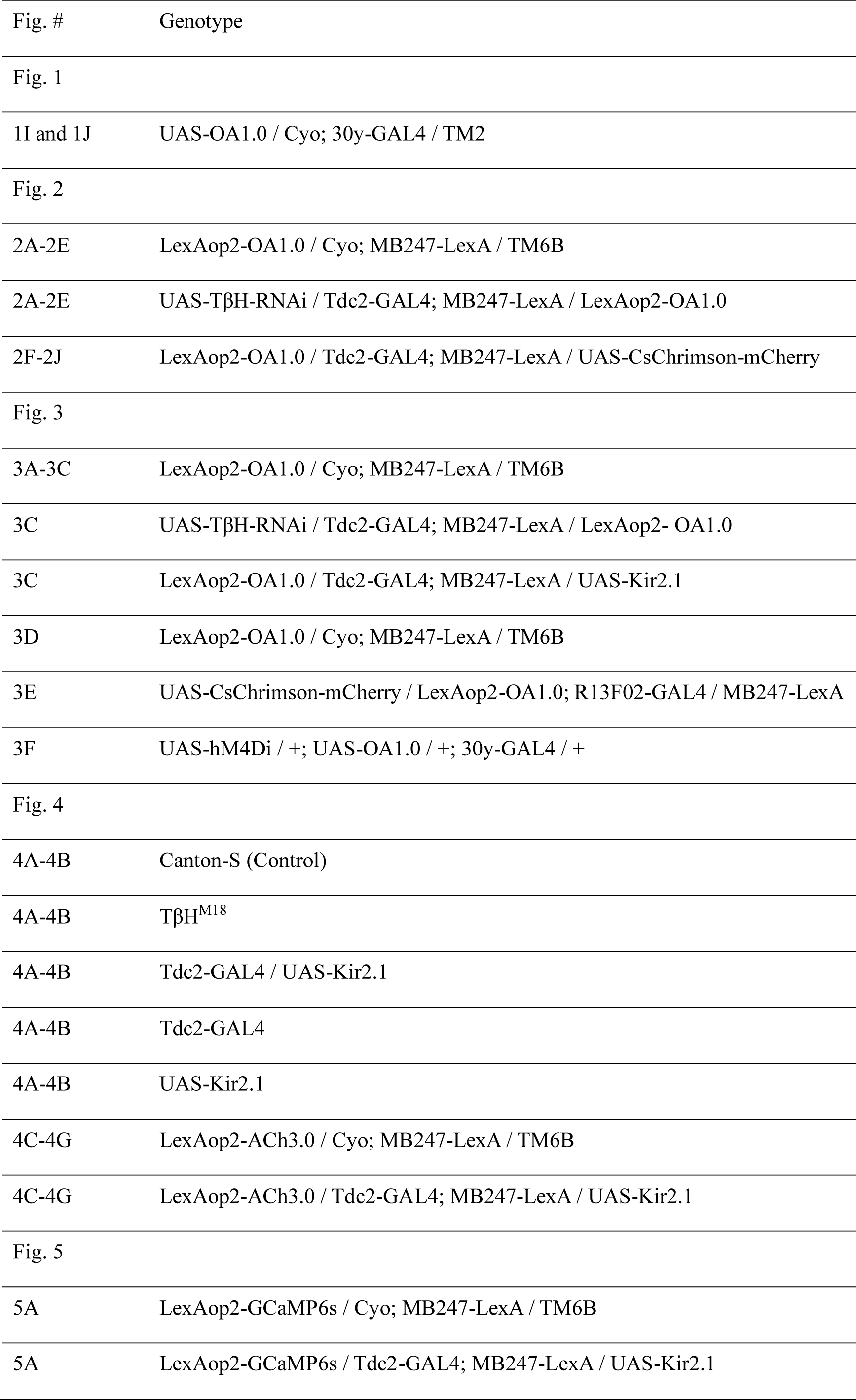
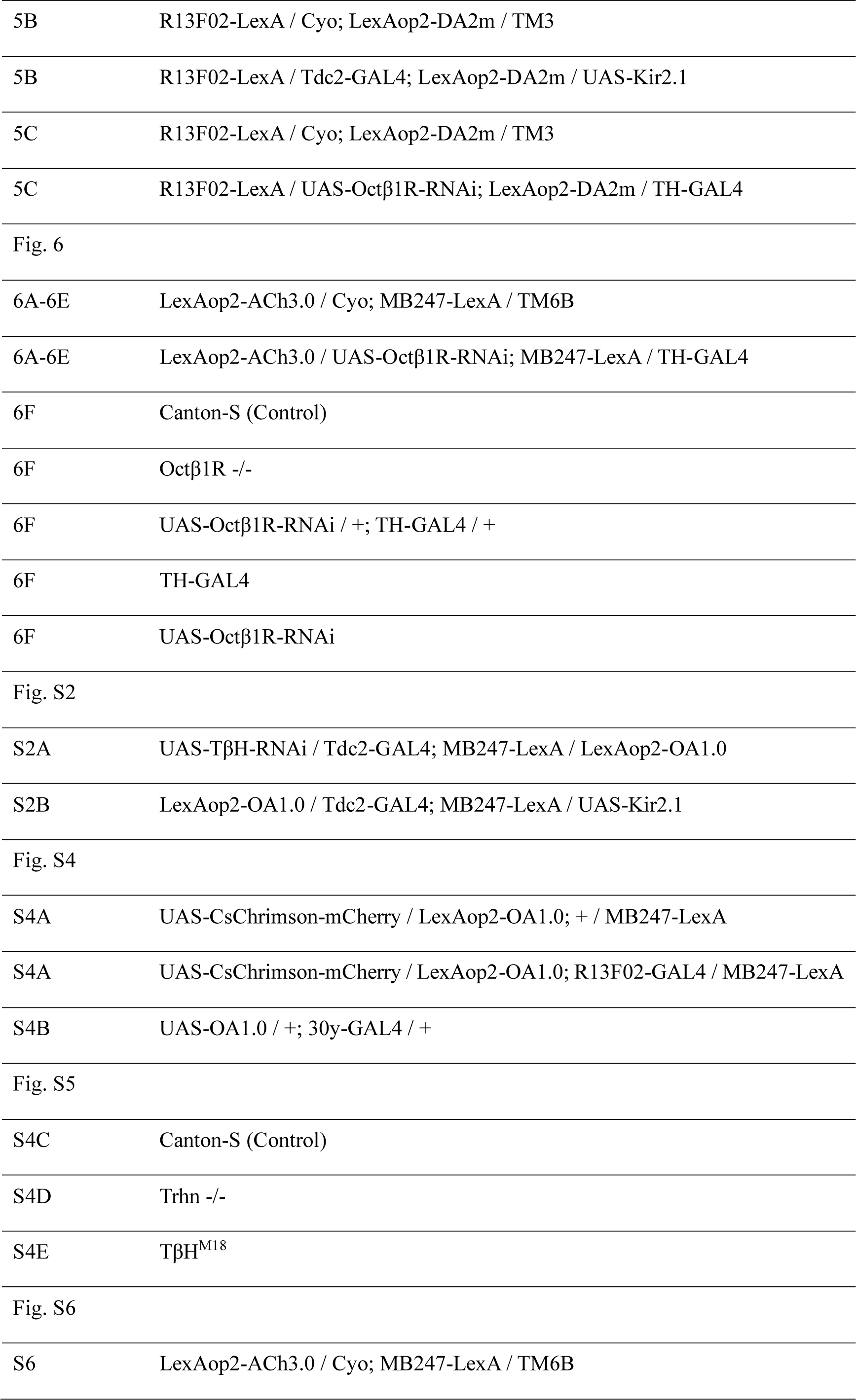

## DETAILED METHODS

### Molecular biology

Expression clones were generated using the Gibson assembly method. PCR was performed to amplify DNA fragments with ∼25-bp overlap using primers (TSINGKE Biological Technology), and T5 exonuclease (New England Biolabs), Phusion DNA polymerase (Thermo Fisher Scientific), and Taq ligase (iCloning) were used to assemble the fragments. Sanger sequencing (TSINGKE Biological Technology) was performed to confirm the plasmid sequences. To characterize the performance of sensors expressed in HEK293T cells, cDNAs encoding the candidate GRAB_OA_ sensors were cloned into the pDisplay vector under the control of the CMV promoter with an upstream IgK leader sequence; a downstream IRES-mCherry-CAAX cassette was included to label the cell membrane and calibrate the sensor’s fluorescence intensity. Spectral properties were measured using plasmids lacking the IRES-mCherry-CAAX cassette. For the Tango assay experiments, genes encoding the WT *Drosophila* Octβ2R and the OA1.0 sensor were cloned into the pTango vector. For the RFlamp cAMP assay, the RFlamp sensor gene was cloned into the pcDNA3.1 vector under the control of the CAG promoter, and OA1.0 and Octβ2R-pHluorin were cloned into the pCMV vector.

### Expression of GRABOA sensors in cultured cells

The GRAB_OA_ sensors were screened and characterized in HEK293T cells, which were grown either in 96-well plates or on 12-mm diameter circular coverslips in 24-well plates. At 60-70% confluency, the cells were transfected using polyethyleneimine (PEI) at a PEI:DNA ratio of 3:1, and experiments were conducted 24-36 hours after transfection. For 1P spectra measurements and Tango experiments, the cells were cultured and transfected in 6-well plates; after transfection, the cells were transferred to either a 384-well or 96-well plate for subsequent experiments.

### Fluorescence imaging of cultured cells

Before imaging, the culture medium was replaced with Tyrode’s solution containing (in mM): 150 NaCl, 4 KCl, 2 MgCl_2_, 2 CaCl_2_, 10 HEPES, and 10 glucose (pH 7.3-7.4). The HEK293T cells grown in 96-well plates were imaged using an Opera Phenix high-content screening system (PerkinElmer), while the cells grown on 12 mm coverslips were imaged using an inverted Ti-E A1 confocal microscope (Nikon). The Opera Phenix high content screening system was equipped with a 20×/0.4-NA objective, a 40×/0.6-NA objective and a 40×/1.15-NA water-immersion objective, a 488-nm laser, and a 561-nm laser; the GRAB_OA_ signal (green fluorescence) was collected using a 525/50-nm emission filter, and the mCherry signal (red fluorescence) was collected using a 600/30-nm emission filter. The fluorescence intensity of the GRAB_OA_ sensor was calibrated using mCherry as the reference. The Nikon confocal microscope was equipped with a 40×/1.35-NA oil-immersion objective and a 488-nm laser; green fluorescence was collected using a 525/50-nm emission filter. To measure the response kinetics, the tip of a glass electrode was placed approximately 10 μm above the cells; this electrode was pulled using a P-97 Flaming/Brown Micropipette Puller (Sutter Instrument) and contained a saturating concentration of agonist or antagonist. A PV800 Pneumatic PicoPump (World Precision Instruments) was used to control the duration of drug delivery. The fast line-scan mode was used to record changes in the local fluorescence signal at the cell membrane, and NIS-Elements software (Nikon) was used to control imaging.

### Tango assay

HTLA cells were cultured in 6-well plates; at ∼70% cell density, the cells were transfected with either wild-type Octβ2R or OA1.0. Twenty-four hours after transfection, the cells were transferred to a 96-well white clear flat-bottom plate, and virous concentrations of OA (ranging from 1 nM to 100 μM) were added to the cells; each concentration was applied in triplicate. The cells were then incubated for ∼16 hours, and the bioluminescent signal was measured. To measure the bioluminescent signal, the culture medium was removed, and 40 μl of Bright-Glo substrate (Promega) was added to the wells. The plate was then incubated at room temperature in the dark for 10 minutes, and the bioluminescent signal was measured using a Victor X5 microplate reader (PerkinElmer). Non-transfected cells were used as negative controls.

### RFlamp cAMP measuring assay

HEK293T cells were transfected with cytoplasmic RFlamp and either the membrane-targeted Octβ2R-pHluorin or OA1.0 sensor. control cells were transfected with only with RFlamp. The cells were then imaged using Operetta CLS (PerkinElmer) before and after the addition of various concentration of OA.

### Spectra measurements

The 1P spectra of OA1.0 were measured using a Safire 2 microplate reader (Tecan). HEK293T cells were transfected with CMV promoter-driven OA1.0 plasmids (with no other fluorescent proteins); after 24 hours, the cells were dissociated with trypsin and transferred to a clear flat-bottom black-walled 384-well plate for measurement. To detect the excitation spectrum, a gradient of 5-nm (20-nm bandwidth) increments of excitation wavelength was applied from 300–525 nm, and the emission wavelength was fixed at 560 nm (20-nm bandwidth). To detect the emission spectrum, a gradient of 5-nm (20-nm bandwidth) increments of emission wavelength was applied from 495–800 nm, and the excitation wavelength was fixed at 455 nm (20-nm bandwidth). The fluorescence values measured at each wavelength in cells transfected with an empty vector were subtracted as background.

The 2P spectra of OA1.0 were measured using a Bruker Ultima Investigator two-photon microscope equipped with a Spectra-Physics InSight X3 laser. The spectra were measured from 700 nm to 1050 nm at 10-nm increments, and the fluorescence values measured in non-transfected cells were subtracted as background.

### Fast-scan cyclic voltammetry (FSCV)

Fast-scan cyclic voltammetry (FSCV) was performed using an ElProScan ELP-3 equipped with an EPC10 USB triple potentiostat (HEKA Electronik GmbH, Lambrecht/Pfalz, Germany). A carbon fiber electrode (7-µm diameter, 100-200-µm length, Tokai Carbon Co., Tokai, Japan) was used as the working electrode, and a KCl-saturated Ag/AgCl microelectrode was used as the reference electrode in the two-electrode configuration. All high-speed voltammograms were recorded using a waveform potential from −0.4 V to +1.1 V at a scan rate of 400 V/s with a 200-ms interval. The carbon fiber microelectrode was held at −0.4 V between scans.

### Two-photon *in vivo* imaging of flies

For the *in vivo* imaging experiments, we used adult female flies within 2 weeks after eclosion. Each fly was mounted onto a customized chamber using tape, and a rectangular section of tape measuring 1 mm x 1 mm above the head was removed. The cuticle between the eyes, air sacs, and fat bodies were carefully removed in sequential order to expose the brain. Throughout the dissection and imaging experiments, the brain was immersed in adult hemolymph-like solution (AHLS) containing (in mM): 108 NaCl, 5 KCl, 5 HEPES, 5 D-trehalose, 5 sucrose, 26 NaHCO_3_, 1 NaH_2_PO_4_, 2 CaCl_2_, and 2 MgCl_2_.

An Olympus FVMPE-RS microscope equipped with a Spectra-Physics InSight X3 dual-output laser was used for the functional imaging experiments. The green fluorescence signals produced by OA1.0, ACh3.0, DA2m, and GCaMP6s were excited using a 920-nm laser and collected through a 495-540-nm filter. The red florescence signals produced by mCherry-tagged CsChrimson were excited using a 1045-nm laser and collected through a 575-630-nm filter.

To apply local electrical stimuli, a glass electrode (with a tip resistance of 0.2 MΩ) was positioned near the horizontal lobe of the MB, and the stimulation voltage was set to 30–50 V. For optogenetic stimulation, a 635-nm laser was used to deliver 1-ms pulses at 10 Hz through optical fibers positioned near the fly’s brain. For odor stimulation, the odorant was initially diluted 200-fold in mineral oil, and air was bubbled through the oil at 200 ml/min, combined with pure air delivered at 800 ml/min, and finally delivered to the fly antenna at 1000 ml/min. 3-octanol (OCT) and 4-methylcyclohexanol (MCH) were used for the experiments in Fig. 4 and 6, and OCT was used for the experiments in Fig. 3 and 5. For all odor-shock pairing experiments, either OCT or MCH was randomly assigned as the CS+, with the other odorant serving as the CS-. For body shock stimulation, two copper wires were attached to the fly’s abdomen, with the voltage set to 80 V. In the experiments shown in Fig. 1I-J and 3D, a small section of the blood-brain-barrier was carefully removed with tweezers before applying the indicated neurotransmitters and compounds.

### Behavioral assay

These experiments were performed in a dark room at 22°C with humidity ranging from 50-60%. Flies that were 24-72 hours old were selected and transferred to a new tube 12 hours prior to the experiment. Prior to training, the flies were placed in the training arm for 2 min to acclimate.

Each train involved ∼100 flies, and the odorants OCT and MCH were diluted to 1:67 and 1:100, respectively, in mineral oil. The odorant-containing mineral oil was delivered to the training and testing arms of a T-maze at a rate of 800 ml/min.

During the training phase, the CS+ odorant was introduced to the airflow for 1 min, followed by 12 electric shocks (the US) delivered at 80 V (1.25 s/pulse) via a copper grid located in the training arm; 45 s after the shocks were applied, the CS- odorant was introduced to the airflow for 1 min.

Following training, the flies were transferred to an elevator and given a 2-min acclimation period before testing. During testing, the CS+ and CS- were delivered from the two ends of the arms for a duration of 2 min, during which the flies were allowed to make their choice. The number of flies in each arm (N) was recorded after testing, and the performance index (PI) of each trial was calculated.

An Arduino microcontroller board was used to synchronize the delivery of various stimuli, including odorants, shock, and 635-nm light.

## QUANTIFICATION AND STATISTICAL ANALYSIS

### Imaging experiments

Images were processed using ImageJ software (National Institutes of Health). The change in fluorescence (ΔF/F_0_) was calculated using the formula [(F-F_0_)/F_0_], where F_0_ is the baseline fluorescence. The signal-to-noise ratio (SNR) was calculated by dividing the peak response by the standard deviation of the baseline fluorescence. The area under the curve was determined using the integral of the fluorescence response (∫ΔF/F_0_).

### Behavioral experiments

The performance index (PI) was calculated using the formula [(N_CS-_ – N_CS+_) / (N_CS+_ + N_CS-_)]. To minimize the potential influence of innate bias, each PI data point was derived using the average of two trials; in one trial OCT served as the CS+, and in the other trial MCH served as the CS+. This average was then calculated using the formula [(PIOCT CS+ + PIMCH CS+) / 2].

### Statistical analysis

Origin 2019 (OriginLab) was used to perform the statistical analyses. Unless otherwise specified, all summary data are presented as the mean ± sem. The paired or unpaired Student’s *t*-test was used to compare two groups, and a one-way analysis of variance (ANOVA) with Tukey’s post hoc test was used to compare more than two groups. All statistical tests were two-tailed, and differences were considered statistically significant at *P* < 0.05. \**P* < 0.05; \*\**P* < 0.01; \*\*\**P* < 0.001; and n.s., not significant.

---

**Figure S1.**
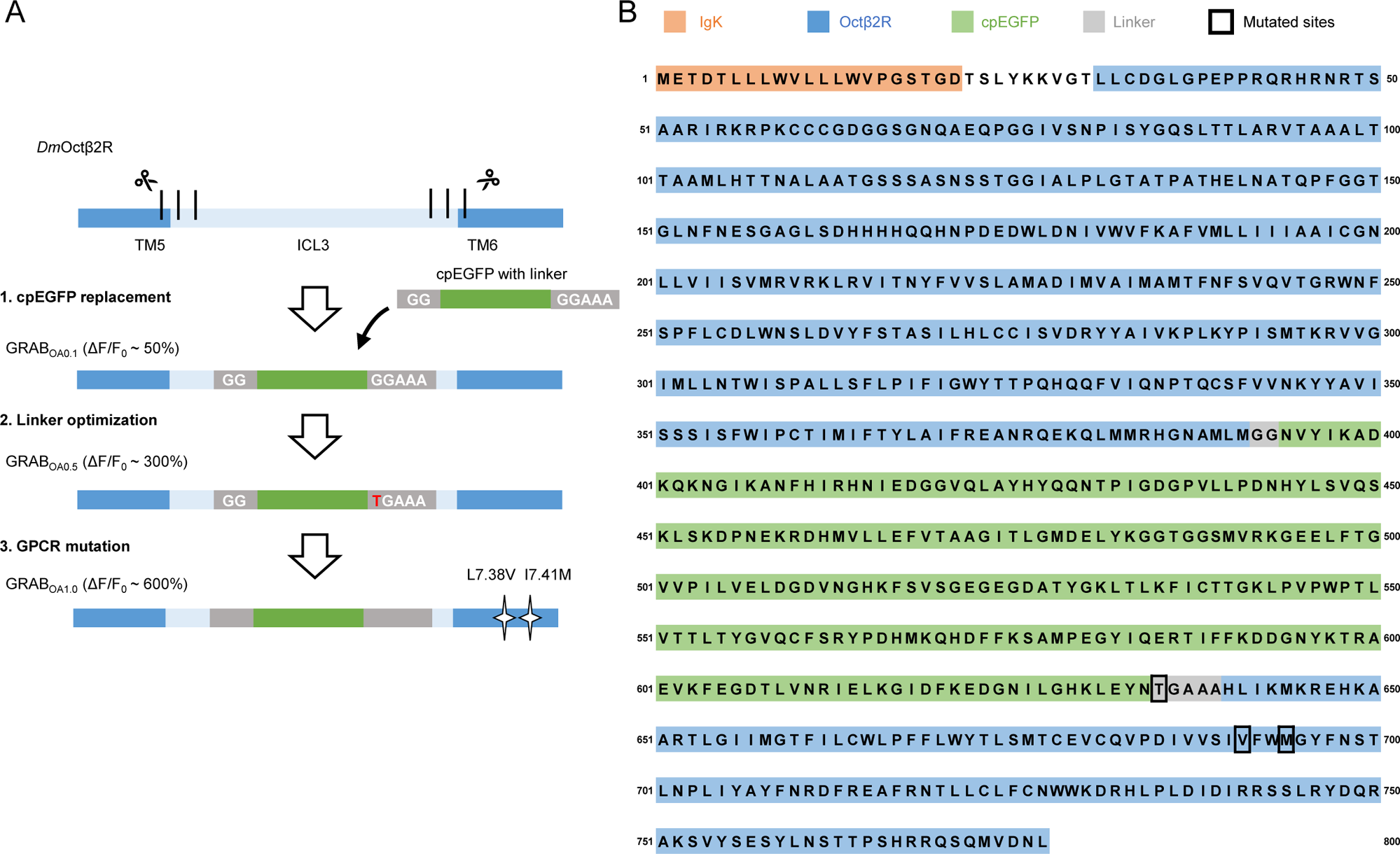
Strategy for designing, optimizing, and screening GRAB_OA_ sensors. (A) Flowchart depicting the process for developing the OA1.0 sensor with a peak response (ΔF/F_0_) of ∼600%. (B) Amino acid sequence of the OA1.0 sensor, with the various domains and mutated sites indicated. Note that the numbering system corresponds to the start of the IgK leader sequence.

**Figure S2.**
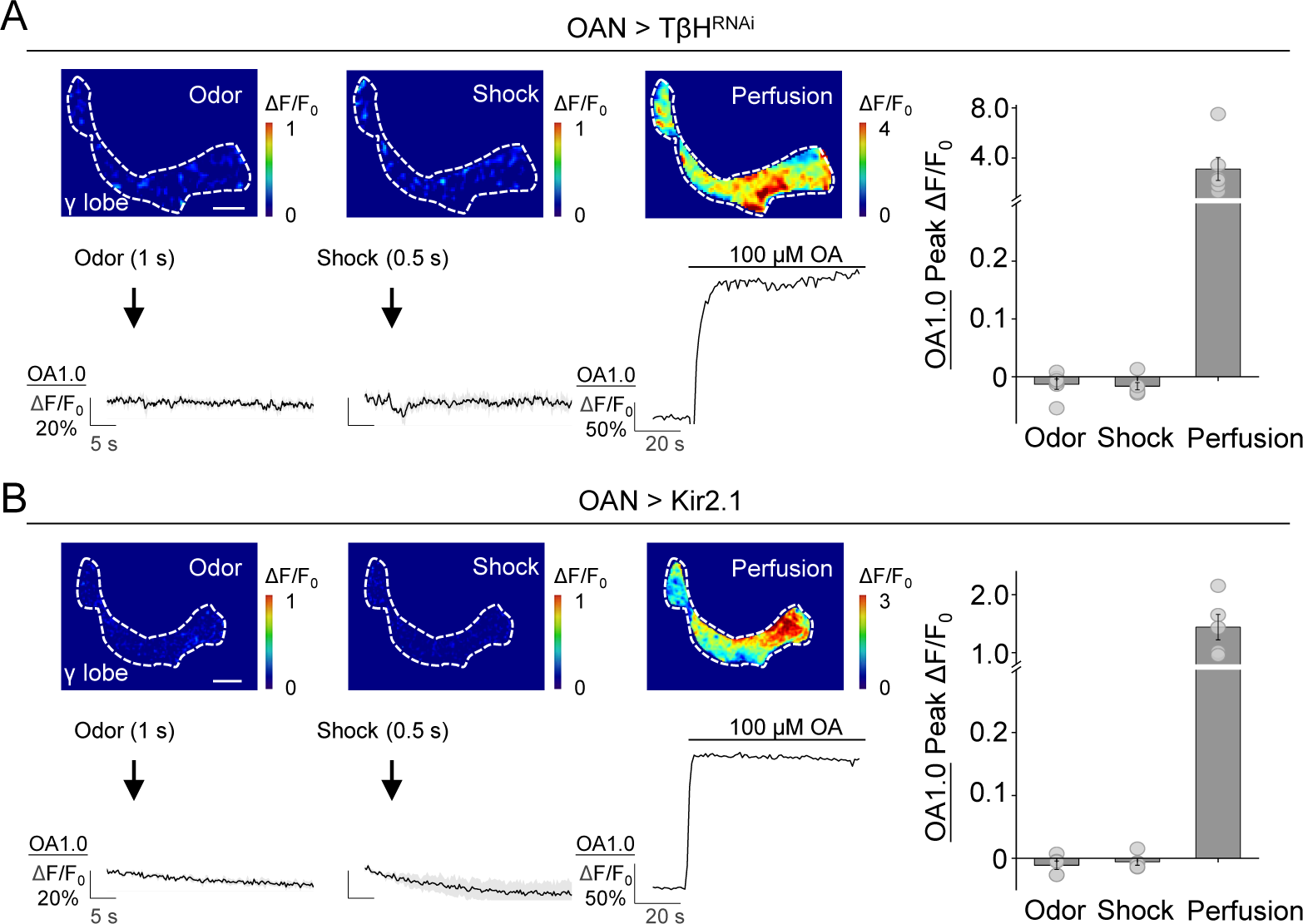
Changes in OA1.0 fluorescence in response to various stimuli measured in OAN > TβH^RNAi^ and OAN > Kir2.1 flies. Representative pseudocolor images (left top), traces (left bottom), and summary (right) of the change in OA1.0 fluorescence measured in response to odor, electrical body shock, and OA perfusion in OAN > TβH^RNAi^ flies (A) and OAN > Kir2.1 flies (B); n = 5-6 flies/group. Scale bars = 20 μm.

**Figure S3.**
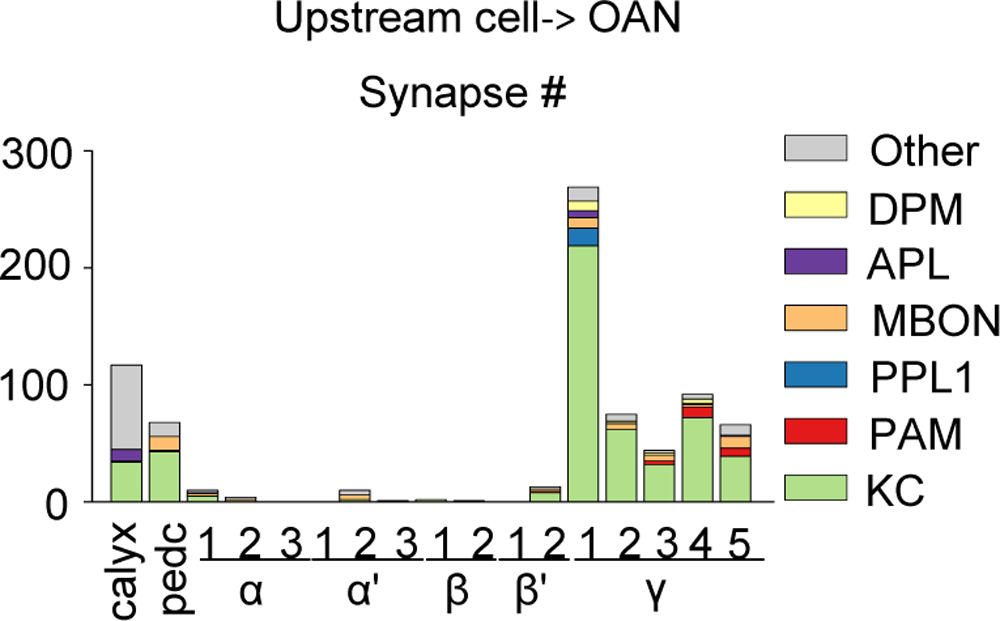
Summary of the number of synapses between OANs and upstream cells in the MB. Pedc, peduncle; DPM, dorsal paired medial; APL, anterior paired lateral neuron; MBON, mushroom body output neuron; PPL1, paired posterior lateral 1 cluster neuron; PAM, protocerebral anterior medial cluster neuron; KC, Kenyon cell; TPM, transcripts per million. Version 1.1 of the hemibrain connectome[56] was used for the analysis, and only synapses with a confidence value >0.75 were included.

**Figure S4.**
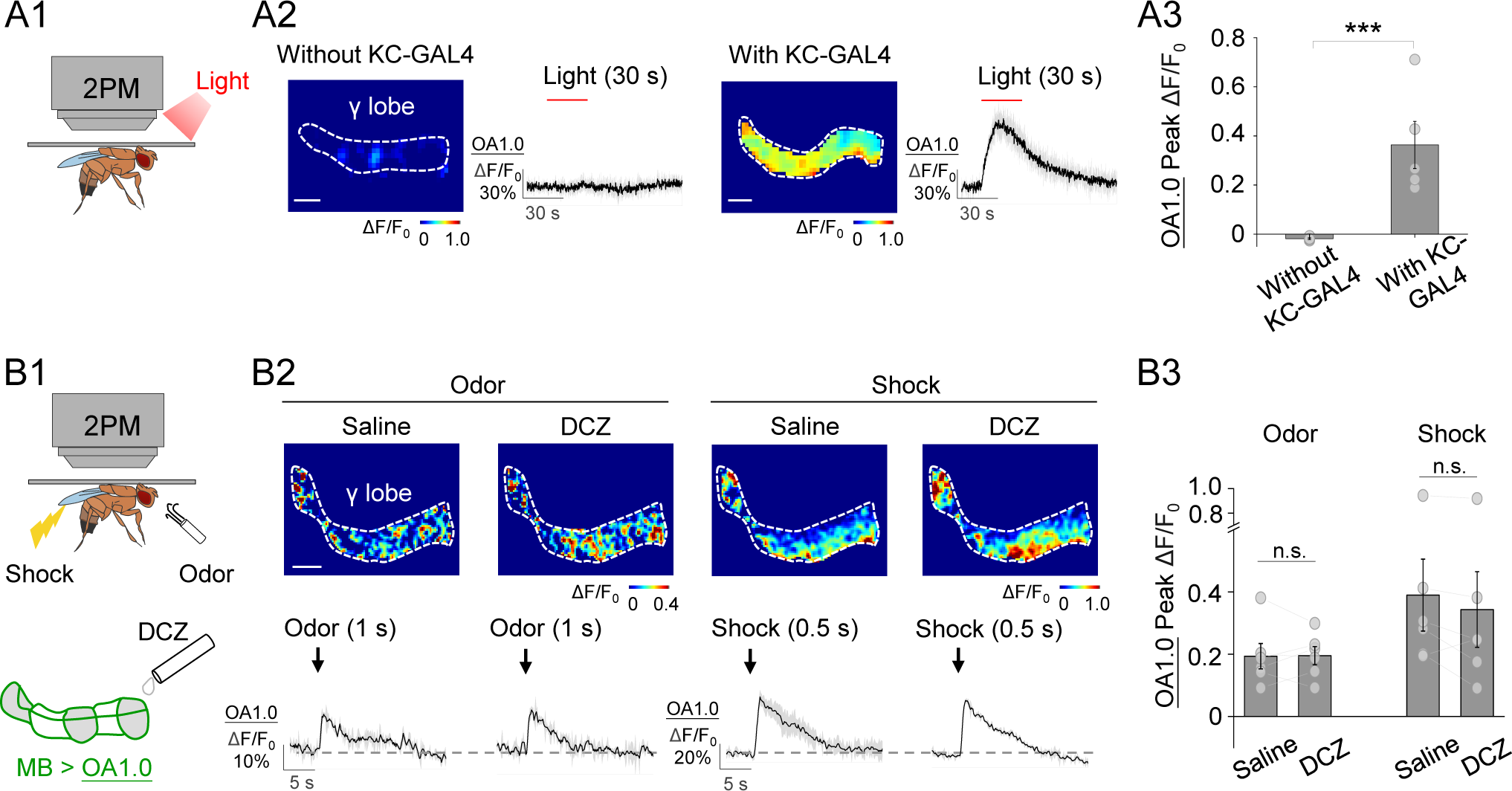
KCs release ACh to trigger OA release, related to Fig. 3. (A) Light stimulation does not lead to OA release in flies with UAS-CsChrimson but without KC-Gal4 driver, ruling out the unspecific effect caused by leaky expression of channelrhodopsin. Shown are schematics (A1) depicting the in vivo imaging setup in which OA was measured with OA1.0 expressed in KCs (MB247-LexA-driven), while the light pulses (1ms/pulse, 635 nm, 10 Hz) were delivered to the brain of the fly only carrying UAS-CsCh-mCherry, but not KC-GAL4. Also shown are representative pseudocolor images, traces (A2), and summary (A3) of the change in OA1.0 fluorescence in response to light pulses (30 s) in flies without or with KC-GAL4; n = 5 flies/group. (B) The hM4Di agonist DCZ does not cause significant effect on odor or shock-evoked OA signals in the γ lobe. Shown are schematics depicting the *in vivo* imaging setup in which OA was measured in the γ lobe using OA1.0 expressed in KCs (30y-GAL4-driven) in the absence or presence of 30 nM DCZ (B1). Also shown are representative pseudocolor images (B2, top), traces (B2, bottom), and summary (B3) of the change in OA1.0 fluorescence in response to odor or electrical body shock in the absence or presence of 30 nM DCZ; n = 6 flies/group. ***p < 0.001, and n.s., not significant (unpaired Student’s t-test). Scale bar= 20 μm.

**Figure S5.**
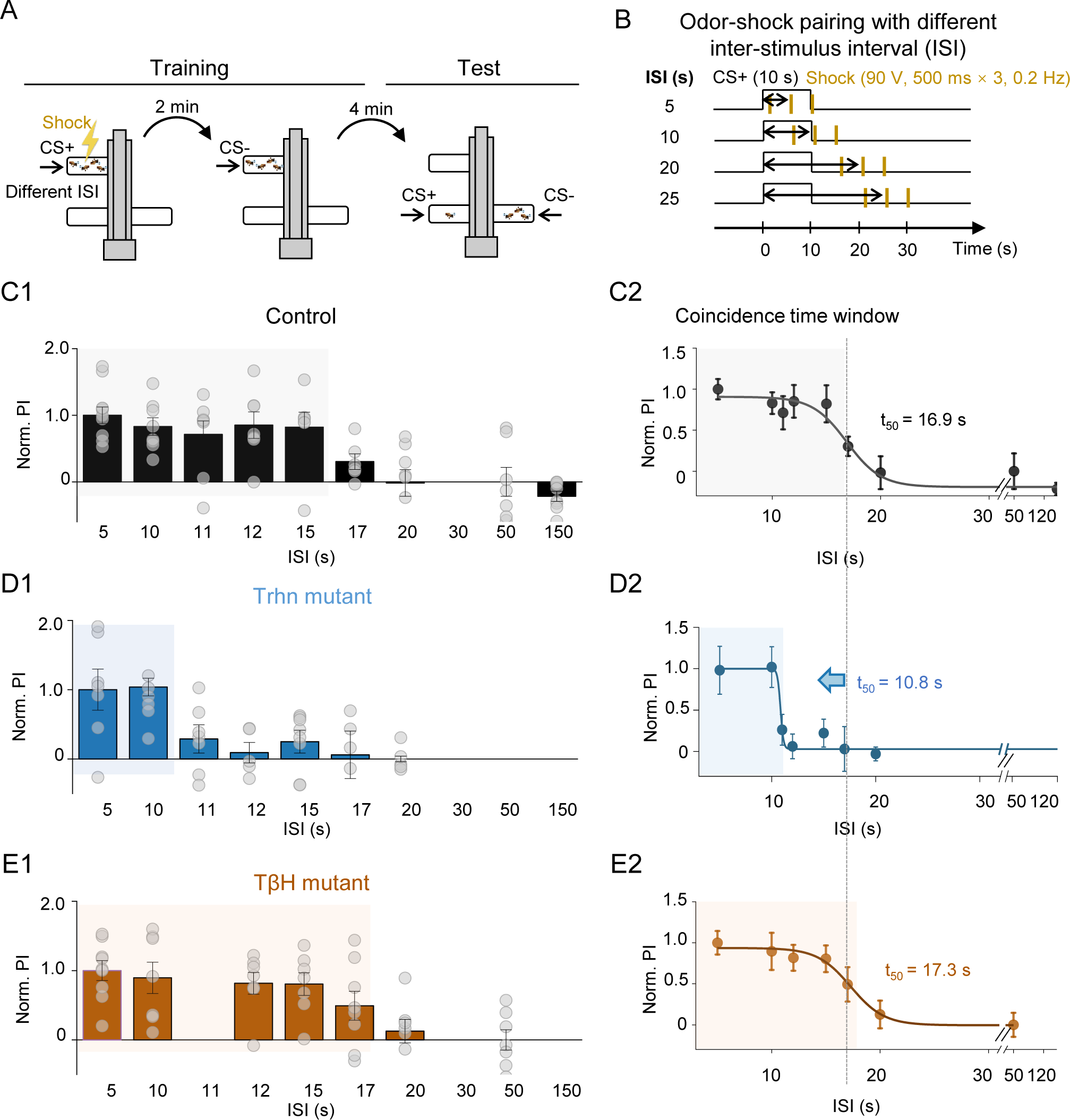
OA signaling does not regulate the coincidence time window of olfactory learning. (A-B) Schematic diagrams depicting odor-shock pairing protocol for measuring how the time interval affects aversive olfactory memory (A) and the effect of varying the inter-stimulus interval (B). (C-E) Summary of the normalized performance index (Norm. PI) measured with the indicated ISI (C1, D1, E1) and normalized PI-ISI profiles fitted to a sigmoid function, with the corresponding t_50_ values shown (C2, D2, E2). The coincidence time window of olfactory learning is defined as the t_50_ for the sigmoid function and is shown as the shaded area. Note that the data presented in panels C and D were reproduced and re-plotted from Zeng et al. (2023)[67]. All group data are presented as mean ± SEM.

**Figure S6.**
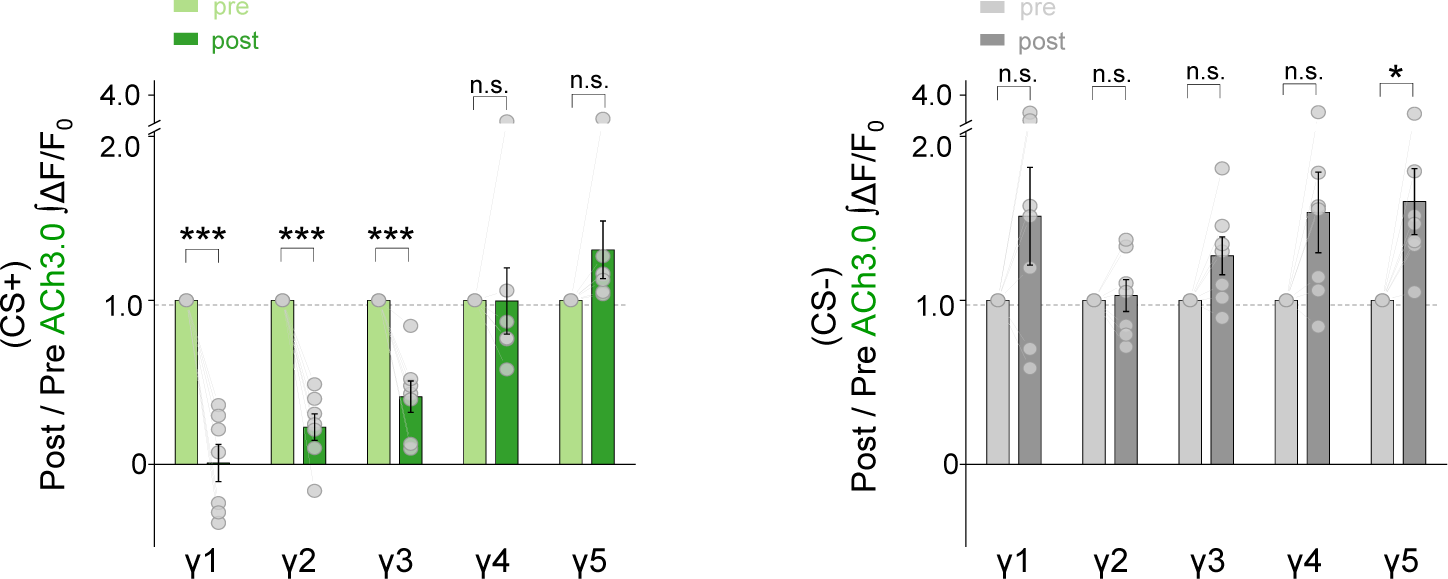
Summary of the relative change in odor-evoked ACh release (post/pre response) following training for the CS+ (left) and CS- (right) measured in wild-type flies. *p < 0.05, ***p < 0.001, and n.s., not significant (unpaired Student’s t-test)

