## Supplementary figures and images for "An octopamine-specific GRAB sensor reveals a monoamine relay circuitry that boosts aversive learning"

### Supplemental figures

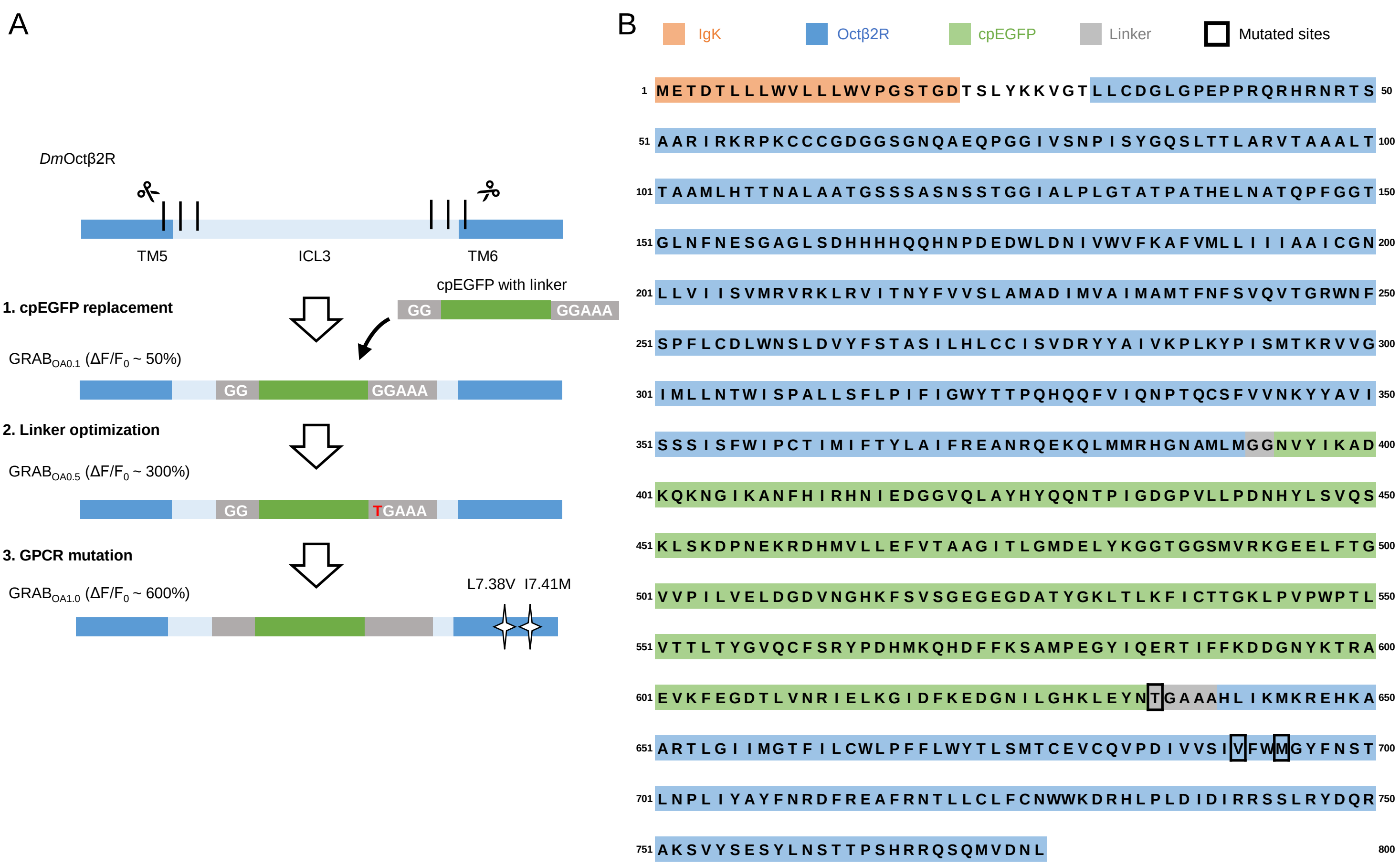

#### Figure S2

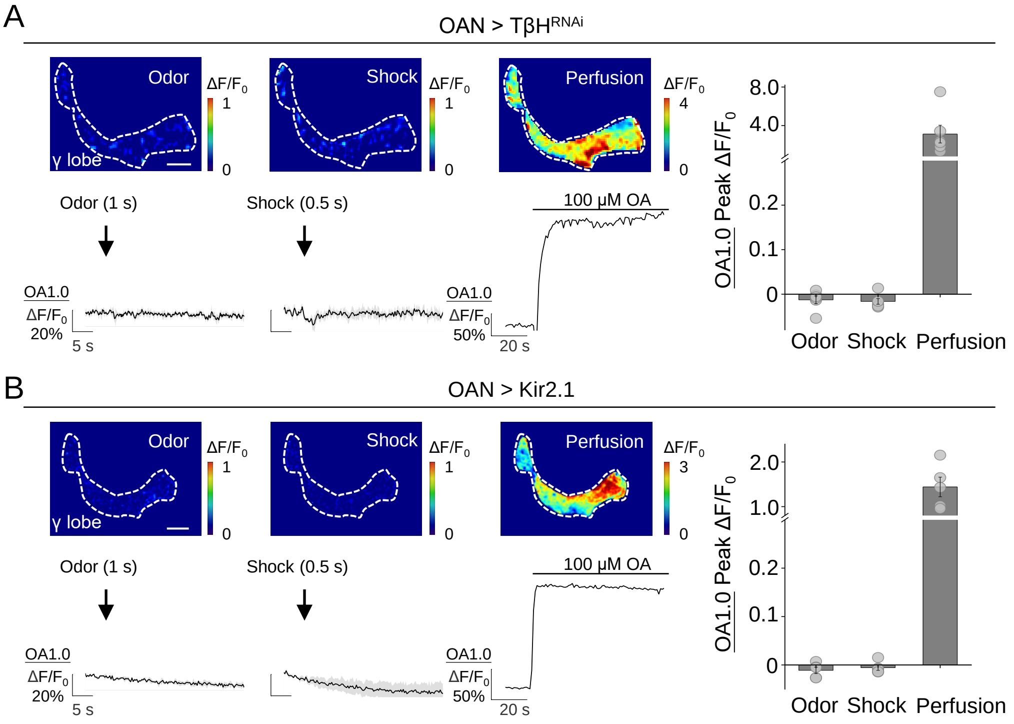


#### Figure S3

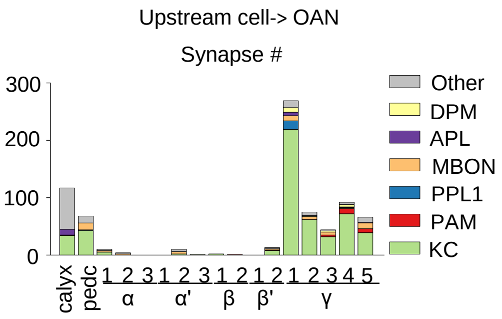

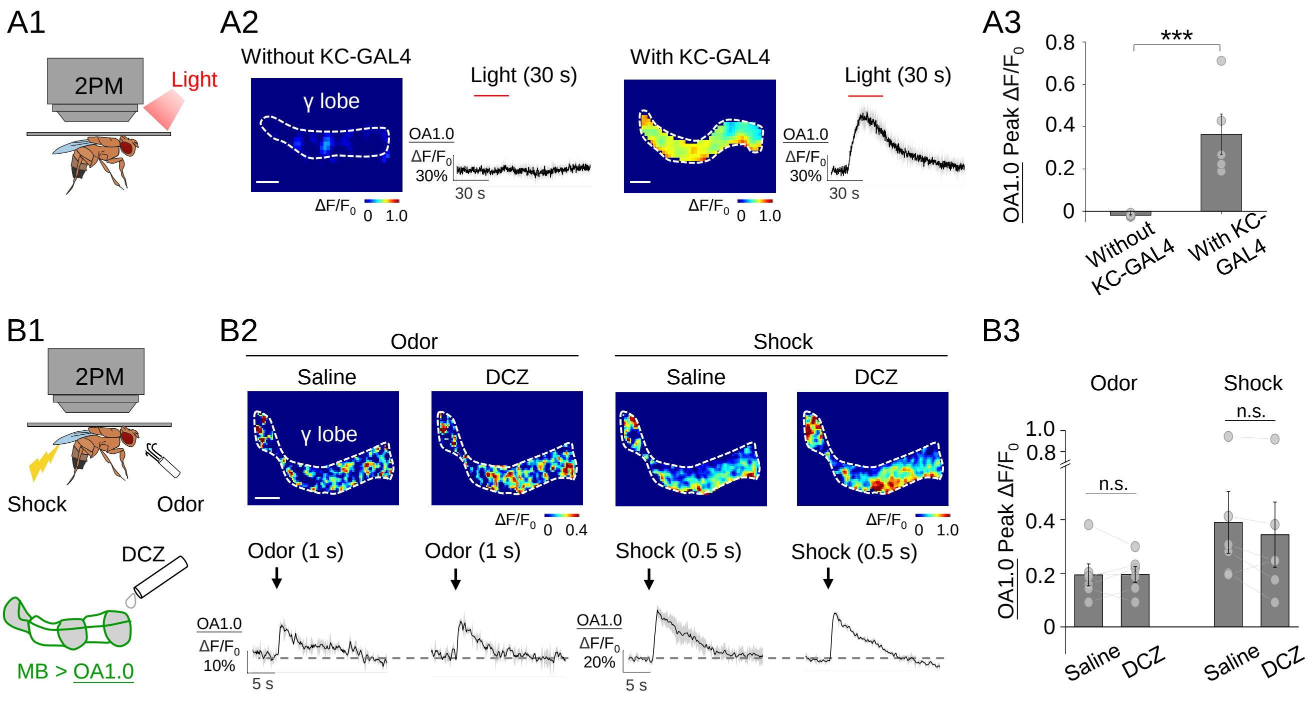

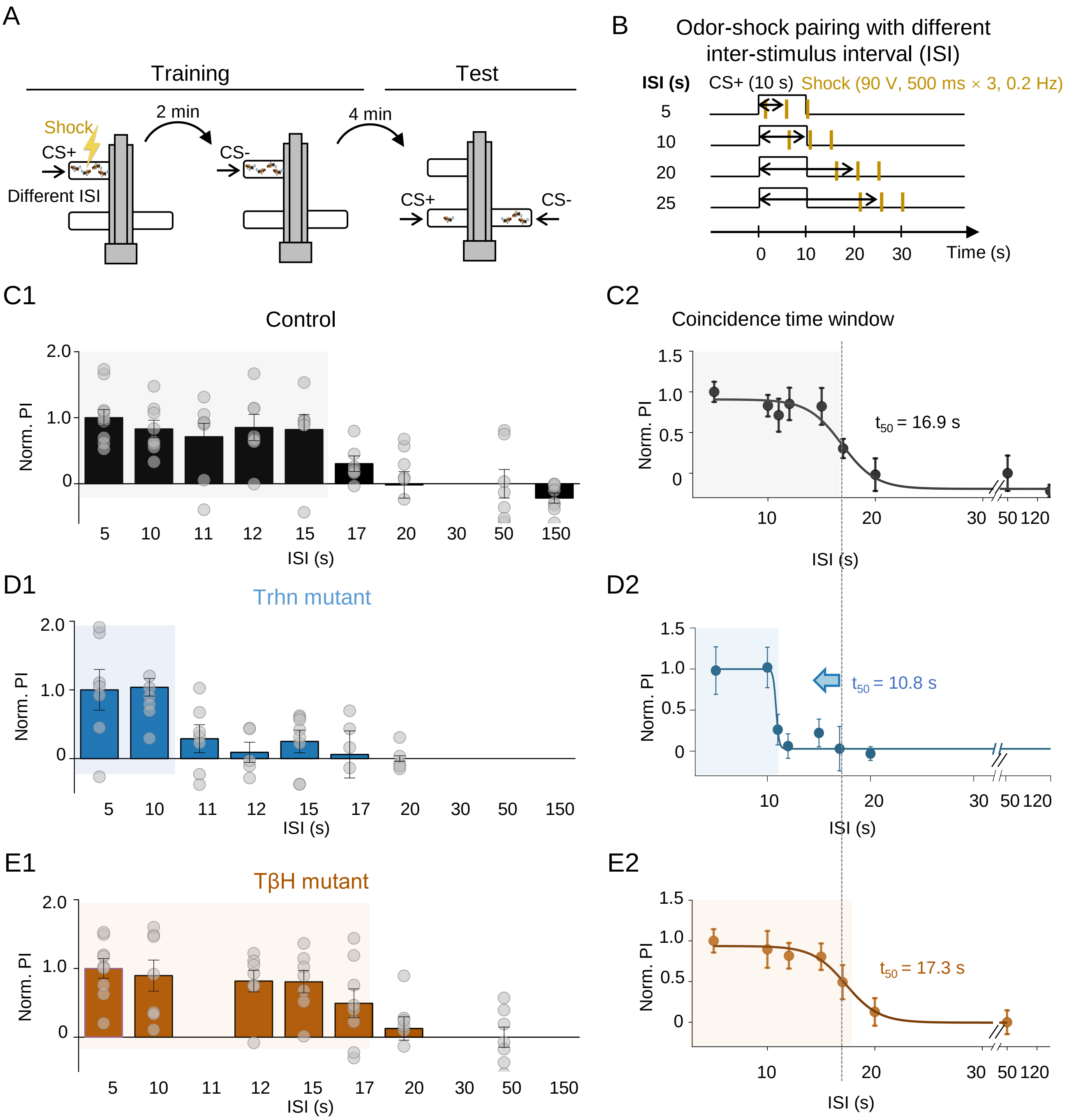

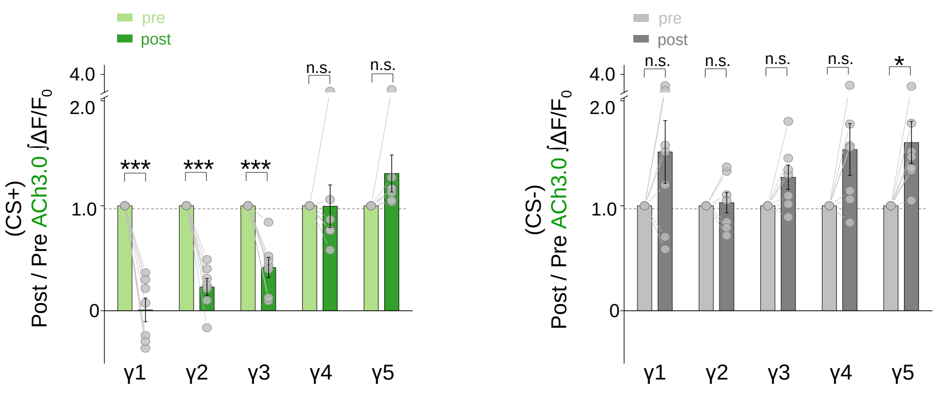
